# Overt and covert prosody are reflected in neurophysiological responses previously attributed to grammatical processing

**DOI:** 10.1101/2020.09.17.301994

**Authors:** Anastasia Glushko, David Poeppel, Karsten Steinhauer

**Affiliations:** Centre for Research on Brain, Language and Music (Montreal, Canada); Department of Psychology, New York University (New York City, NY, USA); Max Planck Institute for Empirical Aesthetics (Frankfurt, Germany); School of Communication Sciences and Disorders, McGill University (Montreal, Canada)

**Keywords:** prosody, cortical tracking, sentence processing, syntax

## Abstract

Recent neurophysiological research suggests that slow cortical activity tracks hierarchical syntactic structure during online sentence processing (e.g., Ding, Melloni, Zhang, Tian, & Poeppel, 2016). Here we tested an alternative hypothesis: electrophysiological activity peaks at sentence constituent frequencies reflect cortical tracking of overt or covert (implicit) prosodic grouping. In three experiments, participants listened to series of sentences while electroencephalography (EEG) was recorded. First, prosodic cues in the sentence materials were neutralized. We found an EEG spectral power peak elicited at a frequency that only ‘tagged’ covert prosodic change, but not any major syntactic constituents. In the second experiment, participants listened to a series of sentences with overt prosodic grouping cues that either aligned or misaligned with the syntactic phrasing in the sentences (initial overt prosody trials). Immediately after each overt prosody trial, participants were presented with a second series of sentences (covert prosody trial) with all overt prosodic cues neutralized and asked to imagine the prosodic contour present in the previous, overt prosody trial. The EEG responses reflected an interactive relationship between syntactic processing and prosodic tracking at the frequencies of syntactic constituents (sentences and phrases): alignment of syntax and prosody boosted EEG responses, whereas their misalignment had an opposite effect. This was true for both overt and covert (imagined) prosody. We conclude that processing of both overt and covert prosody is reflected in the frequency tagged neural responses at sentence constituent frequencies, whereas identifying neural markers that are narrowly reflective of syntactic processing remains difficult and controversial.

## introduction

Language comprehension involves a variety of cognitive mechanisms for processing multiple types of information, from auditory perception to integration of words’ semantic content with the grammatical structure of sentences. While some of these processing mechanisms have parallels across the animal kingdom (ten Cate, 2017), building and processing syntactic structures has been suggested as a unique element of human language that distinguishes it from communication in other animals (Fitch & Hauser, 2004; Berwick, Friederici, Chomsky, & Bolhuis, 2013). From the point of view of syntactic theory (Chomsky, 1959), phrase structure is built from smaller linguistic elements that are combined into increasingly larger units (i.e. from words/morphemes to phrases to sentences), creating a hierarchical structure of grammatical constituents. However, whether this theoretical framework can help describe how the human brain processes language in real time remains controversial (Pylkkänen, 2019). Psycholinguistic and neurolinguistic studies attempting to demonstrate the cognitive processing of hierarchically represented phrasal structures have typically used rather unnatural tasks (such as ‘click’ detection; Fodor & Bever, 1965; Garrett, Bever, & Fodor, 1966) or inferred neurocognitive parsing mechanisms from processing of syntactic errors (e.g., Friederici, Hahne, & Mecklinger, 1996; for review, see Friederici, 2002). This work often produced ambiguous data that could alternatively be explained in terms of semantic or prosodic processing that takes place in parallel to, but is distinct from, syntactic processing (for detailed discussion, see Steinhauer & Drury, 2012). Only few recent studies provided preliminary fMRI and ECoG data on brain responses to syntactic phrase boundaries in grammatical sentences (Pallier, Devauchelle, & Dehaene, 2011; Nelson et al., 2017). The challenges of distinguishing syntactic processing effects per se from those that only appear to be syntactic explain why the recent magnetoencephalographic (MEG) findings by Ding and coauthors (2016) have been widely perceived as a potentially useful new approach to test cortical tracking of hierarchical sentence structures in human listeners (e.g., Friederici et al., 2017; Murphy, 2016). Ding and colleagues’ experiments employed grammatical sentences, used a relatively natural task (listening to connected speech; detecting implausible sentences), and – importantly – explicitly addressed alternative accounts. One such account was a prosodic one: the authors ruled out the contribution of *overt* prosody – i.e., of suprasegmental phonological features including sentence melody and stress patterns (Fery, 2017) – from contributing to their results. Their spoken sentence materials were stripped of tonal pitch and sound intensity changes, and all words within sentences had equal length, ensuring no acoustic cues could mark syntactic boundaries. Despite these precautions, however, it is still possible that Ding and colleagues’ findings might be strongly influenced by *covert* (or implicit) prosody. Covert prosody is known to be activated in absence of any overt acoustic cues, both during silent reading (for review, see Breen, 2014) and in speech perception (Itzhak et al. 2010). The present study was designed to test this hypothesis. Before outlining our specific approach, we briefly summarize Ding and colleagues’ findings and motivate why covert prosody may have played a role in eliciting them. To demonstrate hierarchical syntactic processing, Ding and coauthors (2016) investigated periodic cortical activity using the ‘frequency tagging’ technique. This method can be used to ‘tag’ language characteristics requiring either (a) low-level stimulus-driven (‘bottom-up’) or (b) higher-level cognitively driven (‘top-down’) processing mechanisms. When the authors presented participants with sequences of spoken four-syllable sentences, in which each word consisted of one syllable and lasted exactly 250 milliseconds (making each sentence one second long), they found a robust stimulus-driven 4 Hz rhythm in listeners’ MEG signals. The corresponding 4 Hz peak in the MEG frequency spectrum was found even when English speakers listened to sentences in Mandarin Chinese, demonstrating that this ‘bottom-up’ cortical rhythm simply mirrored the 4 Hz syllable rate of the acoustic signal (envelope tracking), independent of language comprehension (in line with Howard & Poeppel, 2010).

However, when Chinese and English participants listened to 4-syllable sentences in their native language, their MEG signal was characterized by two additional power peaks – at 1 Hz (corresponding to sentence rate) and 2 Hz (corresponding to phrase rate). These two lower frequency effects did not correspond to any acoustic rhythms in the speech signal and must have, therefore, reflected cognitively driven ‘top-down’ brain activity related to understanding and structuring the utterances. In fact, these data were taken to demonstrate the human brain’s ability to track syntactic constituents at multiple distinct levels of the linguistic hierarchy. In both English and Mandarin Chinese, the sentences had been designed such that the first two syllables always created a syntactic noun phrase (NP; e.g., “*new plans*”), while the last two words created a syntactic verb phrase (VP; e.g., “*give hope*”). NP and VP in this “2+2” structure were separated by the sentence’s largest syntactic boundary in mid-sentence position (e.g., “*New plans* | *give hope”*). The authors interpreted the 1 Hz power peak to reflect parsing of the entire sentence (i.e., the largest syntactic constituent), and the phrase-level (2 Hz) peak to represent cortical tracking of the two syntactic units at the next level of the syntactic hierarchy (i.e., NP and VP). This interpretation was supported by an additional condition (in Chinese only) showing that the 2 Hz peak (but not the 1 Hz peak) disappeared when the largest syntactic boundary was placed after the first one-syllable word (“1+3” structure), thus separating two syntactic constituents of unequal length (1 syllable + 3 syllables, as in “*fry* | *to-ma-toes*”). Since all words used in these experiments were recorded separately, had identical length, and their pitch and sound intensity were held constant, Ding and colleagues (2016) provided strong evidence for cortical top-down mechanisms in online speech processing. A computational model links these findings to the larger question of composition in syntactic structures, and the construction of arguments more broadly (Martin & Doumas 2017), lending support to the structure building interpretation.

Covert, implicit prosody processing is one such top-down mechanism, one whose role in the elicitation of neurophysiological power peaks at frequencies of syntactic units remains unknown. Prosodic processing is not limited to bottom-up mechanisms driven by acoustic cues in the speech signal. Instead, readers have been shown to systematically activate covert (implicit) prosodic patterns during silent reading, such as prosodic boundaries that group words into prosodic phrases (Fodor, 1998; Hwang & Schafer, 2009; for review, see Breen, 2014). For example, electroencephalographic (EEG) studies have shown that silent readers reliably elicit a specific brain response for prosodic phrasing (i.e., the Closure Positive Shift), irrespective of whether the phrasing pattern was induced by punctuation (Steinhauer & Friederici, 2001), by long syntactic phrases (Hwang & Steinhauer, 2011), or by an instruction asking participants to imagine certain prosodic pattern while reading (Steinhauer, 2003). Similar prosodic top-down mechanisms have been reported for speech processing as well, especially in the absence of overt prosodic cues (e.g., Itzhak et al., 2010). This top-down prosodic chunking often reflects the reader’s or listener’s initial syntactic analysis, so mentally imposed prosodic phrases may directly correspond to syntactic phrases (Fodor, Nickels, & Schott, 2018; Itzhak et al., 2010; Hwang & Steinhauer, 2011). However, syntax and prosody do not always have a one-to-one mapping, as non-syntactic factors including word length and the symmetry (or balance) of prosodic sister phrases also play a role (Fodor, 1998; de la Crus-Pavía & Elordieta, 2015, Hirose, 2003; Hwang & Schafer, 2009). Phrase length, semantic coherence, and information structure cues can lead to the placement of prosodic breaks at positions where major syntactic breaks are absent (Frazier, Clifton, & Carlson, 2004; see also discussion of such instances in Samek-Lodovici, 2005; Wagner & Watson, 2010; Shattuck-Hufnagel & Turk, 1996). It is fair to assume that covert prosodic phrasing patterns will reflect the high variability of prosodic realizations seen in speech production with many prosodic boundaries being only optional (Allbritton, McKoon, & Ratcliff, 1996; Schafer, Speer, Warren, & White, 2000) and being inserted, for instance, driven by individual working memory capabilities (Swets, Desmet, Hambrick, & Ferreira, 2007).

Thus, a given syntactic structure is often compatible with multiple distinct prosodic groupings, and a given prosodic structure may be applicable to multiple syntactic structures. For example, the sentence “[*John*]_**NP**_ [*likes* [*big trees*]_NP_]_**VP**_” has a 1+3 syntactic structure, where the subject NP *John* is followed by a 3-word VP (consisting of the verb *likes* and the object NP *big trees*). Prosodically, however, a 2+2 grouping (*John likes* | *big trees*) would be perfectly acceptable. Applied to Ding and colleagues’ materials, these considerations point to a confound between syntax and covert prosody. Their 2+2 syntactic structure is compatible with a 2+2 prosodic structure with a prosodic boundary in mid-sentence position. In contrast, their 1+3 syntactic structure consisting of a monosyllabic verb and a trisyllabic object NP (“*fry to-ma-to”*, or *“drink Oo-long tea”*) is incompatible with a mid-sentence prosodic boundary, because it would separate syllables belonging to the same word (“*fry to* | *ma-to”; “drink Oo* | *long tea”*). Thus, in both structures, the only possible prosodic grouping is identical to the syntactic phrasing. In addition, the entire 4-word utterance in all cases would correspond to the largest prosodic group (a so-called ‘intonational phrase’), which would provide a prosodic account for the sentence-level 1 Hz peak as well. The notion of covert prosody becomes especially relevant when it comes to frequency tagging studies, where sentences are typically presented in a blocked design, such that a given trial of 12 sentences contains either only 2+2 sentences or only 1+3 sentences. This way, listeners could quickly develop a covert prosodic template during the first few sentences and then apply this template to the remaining sentences of a trial, thereby eliciting the 2 Hz peak in 2+2 but not in 1+3 sentences. The initial motivation for generating these prosodic groupings can either be syntactic or non-syntactic in nature, but lexical rather than syntactic reasons seem to prevent a 2+2 prosodic grouping in 1+3 syntactic structures used by Ding and colleagues.

Given the potential confounds between syntactic and covert prosodic phrasing in Ding and coauthors’ materials, it is not unlikely that their sentence (1 Hz) and ½ sentence (2 Hz) MEG power peaks do not reflect the hierarchical levels of syntactic structure, but rather the prosodic grouping of words.

### Present study

To test the hypothesis that covert prosody may have contributed to the 1 Hz (sentence-level) and 2 Hz (½ sentence, phrasal frequency) peaks attributed to syntactic processing, we present EEG experiments with German sentence materials that unconfounded syntactic and prosodic phrasing. Similar to Ding and colleagues (2016), we created 2+2 and 1+3 syntactic structures. However, in contrast to their materials, our 1+3 Syntax condition was still compatible with a 2+2 prosodic grouping (similar to the sentence example provided above, i.e., *John likes big trees*). In our first “No Prosody” experiment, we adopted Ding and coauthors’ (2016) paradigm and presented series of 4-word sentences without any prosodic information. We predicted that if syntax alone was responsible for the sentence- and phrase-level peaks, the 1+3 Syntax condition should replicate the original findings and not elicit the ½ sentence frequency peak (see Figure 1a). However, if covert prosody was involved, this condition should now elicit both the sentence and the ½ sentence peaks (Figures 1c and 1e). Further, in our “Prosody” experiment, we created prosodic contours that were expected to differentially interact with the two syntactic structures (2+2 and 1+3, respectively; for an illustration, see Figure 1). These contours were applied to the sentences from the “No Prosody” experiment either overtly, by modulating the auditory sentence materials, or covertly, by asking participants to imagine a specific prosodic contour while listening to sentences without overt prosodic cues. Our expectation was that *both overt and covert* prosody should increase the ½ sentence frequency peak in sentences with a 2+2 syntactic structure, but not in those with a 1+3 structure. Finally, task effects and their interaction with prosodic processing were studied in an additional “No Semantic Task” experiments (see Supplementary Materials *C*).

**Figure 1.**
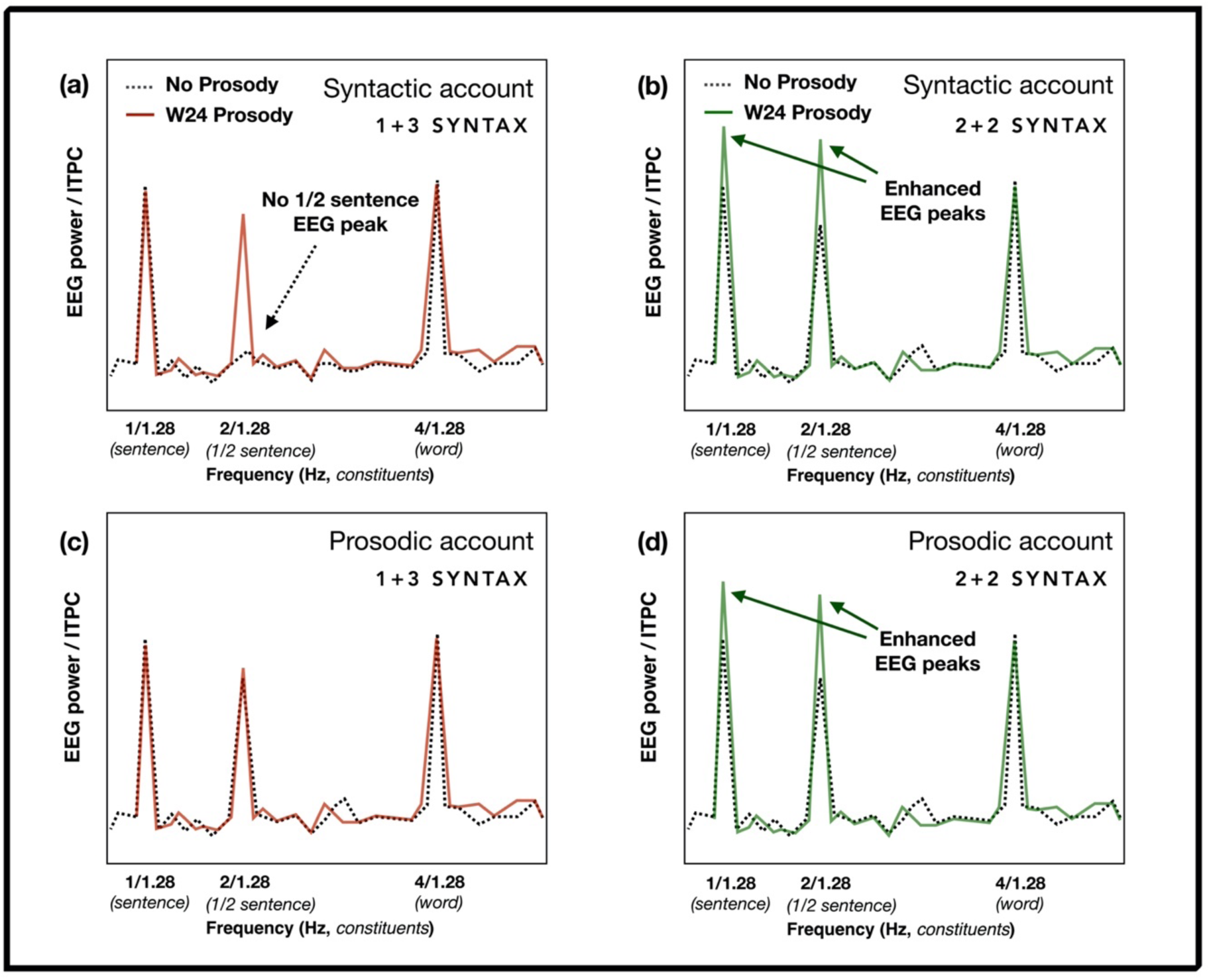
Predictions for No Prosody and Prosody experiment outcomes based on different theoretical assumptions. The top row represents the syntactic account of Ding et al. (2016) findings (panels *a* and *b*), while the predictions driven by the prosodic account can be seen in the bottom row (panels *c* and *d*). If covert prosody does not play a role in the elicitation of EEG power peaks at syntactic constituent frequencies when overt prosodic cues are neutralized (i.e., No Prosody condition), no ½ sentence peak is expected for the 1+3 Syntax condition (dotted line in panel *a*). In contrast, in the 2+2 Syntax condition, this peak would be present (dotted line, panel *b*). Alternatively, if the ½ sentence peak can be accounted for by prosody, both syntactic structures would elicit it in the No Prosody conditions (panels *c*-*d*). In the Prosody conditions (i.e., W24 Prosody), we predicted, independently of the account, to see an interaction between syntactic and prosodic structures: when syntax and prosody are aligned, we expected to see enhanced EEG responses at sentence constituent frequencies (panels *b* and *d*). When they are misaligned, this effect would be significantly weaker or non-existent (panels *a* and *c*).

## Methods

### Participants

Twenty-six participants (age range: 19-45 years, mean age = 27, age SD = 6; 15 females, 10 males) took part in both the No Prosody and Prosody experiments. All participants were recruited and tested at McGill University in Montreal, most of them visiting Canada for work-and-travel purposes. They had acquired German language from birth and considered it their dominant language. The inclusion criteria for the study were the absence of neurologic or psychiatric disorders and hearing impairments, as well as normal or corrected vision. Participants provided written informed consent and received monetary compensation ($20/hour) for their time.

We assessed handedness using the Edinburgh Handedness Inventory ensuring all participants were right-handed (Oldfield, 1971). Participants filled out detailed in-house questionnaires about their language background and musical expertise ensuring all of them were native speakers of German and non-musicians. All parts of the study were approved by McGill’s Faculty of Medicine *Institutional Review Board* (IRB) prior to data collection.

### Materials

#### Speech synthesis

The four-word German sentences used in the experiment were synthesized word-by-word with a built-in Apple synthesizer (the Anna voice). All words were monosyllabic, and their speech signals were exactly 320 ms long. The pitch of each word (and thus of the entire sentence) was flattened, and the intensity was normalized to 70 dB in Praat (Boersma & Weenink, 2019). The words were concatenated into 80 semantically plausible and 24 semantically implausible 4-word sentences, which were further concatenated into *trials* each comprising 12 sentences (48 words). The semantically implausible ‘outlier’ sentences were arranged by re-combining words from two semantically plausible sentences (e.g., *Das Zelt lacht lahm*; lit.: “The tent laughs lamely”) and were used as targets in the outlier detection task (see below). Each sentence was repeated 8 to 9 times within the same experimental block but never within the same trial. Each trial lasted for 15.36 seconds (12 sentences x 4 words x 320 ms; no pauses were introduced between words, phrases, and sentences), identical to the trials in Ding and colleagues (2017). For the two types of syntactic structure and for each type of prosodic contour used in the study (see below), we created 22 trials without any implausible outliers. In the experimental conditions that employed the outlier detection task, we added 8 additional trials with one outlier sentence each, but these were not subjected to subsequent data analysis.

#### Syntactic structure of the sentences

Sentences followed one of the two types of syntactic structures. In the case of the *2*+*2 Syntax* (40 sentences), sentences consisted of two syntactic phrases of equal length. The first phrase was a noun phrase (NP), consisting of a determiner and a noun, while the second one was a verb phrase (VP), most frequently comprised of a verb and an adverb (e.g., *Der Tisch steht da*; lit.: “The table stands there”, or “The table is over there”). In rare cases, the verb phrase (VP) consisted of a particle verb with the corresponding particle replacing the adverb (e.g., *Mein Boot kippt um*; English: “My boat tips over”). In the *1*+*3 Syntax* (40 sentences), the first phrase in each sentence included a one-word NP (i.e., a name), and the second phrase was represented by a 3-word VP (typically, a verb and its complement, e.g., a verb + a determiner/preposition + a noun; e.g., *Lars mag das Bild*; English: “Lars likes the picture”; see Supplementary Materials *A* for the full list of sentences and additional details on their characteristics). The two types of syntactic structures were compared. Given that the current study used EEG (and not MEG, like the only frequency tagging study using 1+3 Syntax constructions with phrases of non-equal length), we ran a control experiment in a separate group of participants to establish that the EEG effects for 1+3 groupings are analogous to the ones reported in Ding and coauthors’ (2016) study (see Supplementary Materials *B*). Our results confirmed that 1+3 rhythm elicits an EEG spectrum similar to the MEG spectrum reported by Ding and colleagues (2016) and can, therefore, be contrasted with the 2+2 Syntax sentences in our main study. For both types of syntactic structure, the (acoustically unmarked) sentence boundary appeared once every 1.28 seconds (after four words) at a frequency of 1/1.28 (0.78) Hz (*sentence frequency*), and single words appeared every 320 ms (i.e., at a *word* frequency of 3.125 Hz). However, only in the 2+2 Syntax sentences (where the phrase boundary between the NP and the VP occurred after two words), syntactic phrases were isochronous and appeared at a constant frequency of 1.56 Hz (every 640 ms), that is, at *½ sentence frequency*.

#### Prosodic manipulations of the sentences

The sentences concatenated from words with neutralized prosody as described above constituted the *No Prosody* condition (henceforth, *NoP;* used in the No Prosody experiment) that was to be contrasted with the data from the *Overt* and *Covert Prosody* conditions (henceforth, *OvP* and *CovP* respectively; used in the Prosody experiment). As the general idea of our prosodic manipulation was to create prosodic patterns that would selectively support one syntactic structure (e.g., 2+2) while conflicting with the other one (e.g., 1+3), the most straightforward acoustic manipulation would have been to either insert pauses at a boundary position or increase the duration of pre-boundary syllables. This kind of prosodic manipulation changes the duration of pre-boundary words and has not only been found to be the most reliable boundary marker in natural speech, but has also been successfully used in previous studies to create cooperative and conflicting syntax-prosody pairings (e.g., Kjelgaard & Speer, 1999), including in EEG studies (Steinhauer et al., 1999; Bögels, Schriefers, Vonk, Chwilla & Kerkhofs, 2009; Pauker, Itzhak, Baum, & Steinhauer, 2011). However, in a frequency tagging study that crucially depends on the invariable duration of all syllables, phrases, and sentences (see above and Ding et al., 2016), durational manipulations are not an option. Instead, we manipulated pitch and intensity, two prosodic dimensions that also contribute to prosodic boundary marking (Streeter, 1978; Beckman, 1996; Männel, Schipke, & Friederici, 2013; Roll, Horne, & Lindgren, 2010). To this end, we synthesized artificial pitch and sound intensity contours in Matlab R2019a (Mathworks, 2011). The resulting artificial prosodic contours were then imposed onto the sentences of the No Prosody condition (using Praat, Boersma & Weenink, 2019), thereby creating the Overt Prosody condition (see Figure 2).

**Figure 2.**
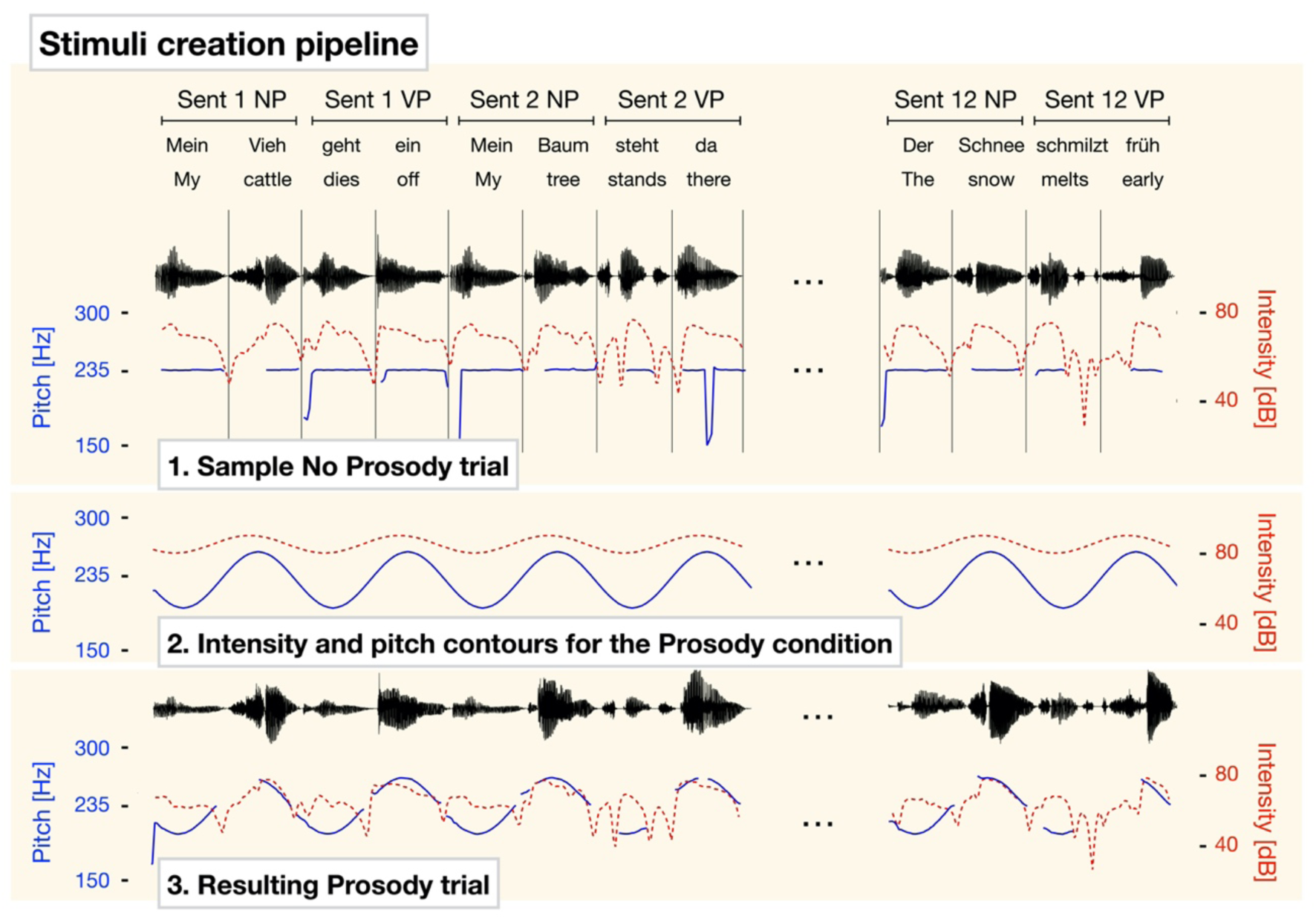
Stimulus development scheme. Single words were synthesized and concatenated into trials (12 sentences each; sample sentences are taken from one of the 2+2 Syntax trials). In the No Prosody experiment (1), words were synthesized, and prosodic cues were neutralized: i.e., there were no pauses between words within trials, all words were 320 ms long, pitch was flattened, and sound intensity was constant across words. Artificial prosodic contours were then created using 1.56 Hz (½ sentence rate) sine waves (2) and imposed with neutralized prosody to create stimuli for the overt prosody trials (to be used in the Prosody and the No Semantic Task experiments). Pitch contour is depicted in blue (note that infrequent sudden drops of pitch values typically reflect unavailability of pitch information due to unvoiced phonemes), and sound intensity is represented by red lines. (3). Audio files for all stimuli are available upon request.

In the Overt Prosody condition, the maxima of sound intensity and pitch were placed on Words 2 and 4 (hence, W24 contour). That is, the fluctuations of pitch and intensity appeared at the ½ sentence frequency (1.56 Hz). Avoiding strategic carry-over effects across sentence conditions, other prosodic contours were tested in alternation with the W24 contour, but these are irrelevant for our present findings and will be reported elsewhere. We imposed the W24 contour onto all experimental sentences of both 2+2 and 1+3 Syntax structures. We considered the W24 prosodic contour to be aligned with the syntactic phrasing of the 2+2 Syntax sentences: using prosodic boundary cues suitable for a frequency tagging paradigm (i.e., avoiding changes in lengthening), we created a simplified model of a prosodic boundary placed at words wrapping up the two syntactic phrases. We predicted an enhancement of ½ sentence rate EEG responses in the case of syntax-prosody alignment (2+2 Syntax) and expected this effect to be stronger than any analogous effect in the 1+3 Syntax sentences. This is because the 1+3 Syntax sentences with the W24 prosodic contour present the case of syntax-prosody misalignment: prosodic changes are not placed at the phrase-final position (i.e., the second word in 1+3 Syntax sentences does not wrap up a syntactic phrase) and syntactic phrases are not repeated at the frequency of ½ sentence, at which the prosodic changes occur. In this case, we expected to see reduced brain activity at the frequency of the sentence due to participants hindered ability to syntactically chunk sentences. That is, we predicted that prosodic and syntactic effects would be non-additive and that prosodic and syntactic analyses would interact during online sentence processing.

We tested all experimental sentences for intelligibility in the Overt Prosody conditions in 7 pilot participants who did not subsequently participate in the EEG recordings, while the No Prosody intelligibility data were collected from every participant at the beginning of the main EEG experiment (including 11 participants who did not go through the full versions of the No Prosody and the Prosody experiments; see Supplementary Materials A).

In the CovP condition, the No Prosody sentences were preceded by corresponding OvP sentences (see Procedure below). Contrasting the OvP and the CovP conditions allowed us to identify the role of ‘overt’ prosody (acoustically realized in OvP sentences) and imagined ‘covert’ prosody (in CovP sentences).

### Procedure

Every participant visited the lab for 5-6 hours, including a 3.5-4.0 hour period of EEG recording involving three experiments with multiple breaks throughout the EEG session. During the EEG cap setup, participants filled out behavioural questionnaires. After that, they performed a stimulus familiarization task. The experimenter explained to the participants that the stimuli were synthesized and the speech rate was relatively high, which is why some of the sentences might possibly be difficult to understand right away. To avoid any comprehension problems during the EEG study, participants had an opportunity to read through the full list of sentences (including the semantic outliers) prior to the experiment and then performed a computerized sentence intelligibility task (note that exposing participants to the stimuli prior to the main experiment was done in previous research as well; Jin et al., 2018). In this task, participants listened to every sentence (with a maximum of two replays) and typed in what they heard. Using this task, we were able to verify that all participants understood the vast majority of the sentences: on average, they correctly typed in 100 out of 104 sentences (for results, see Supplementary Materials A). Following the behavioural task, the main series of EEG experiments started.

We conducted three experiments (see Figure 3 for experimental structure). Every participant started with the No Prosody experiment that served to establish a baseline for (syntax and, potentially, default covert prosody) processing 1+3 and 2+2 Syntax sentences. Next participants engaged in the No Semantic Task (see below) and the Prosody experiments (whose order was counter-balanced across participants). At the end of the study, we repeated the No Prosody experiment (with a randomized trial order different from the one at the beginning of the experiment) to control for the familiarity with the sentences between the No Prosody and the Prosody experiments and participants’ fatigue. Trials in the Prosody experiment were presented in blocks containing trials with the same prosodic and syntactic structure. The order of the blocks was counter-balanced across participants. The data from the two runs of the No Prosody experiment were averaged after ensuring the main patterns were unaffected by whether the data were recorded at the beginning or at the end of the experiment (see Results). This order allowed for minimal influence of the prosodic contours from the Prosody experiment on the processing of sentences in the No Prosody experiment.

**Figure 3.**
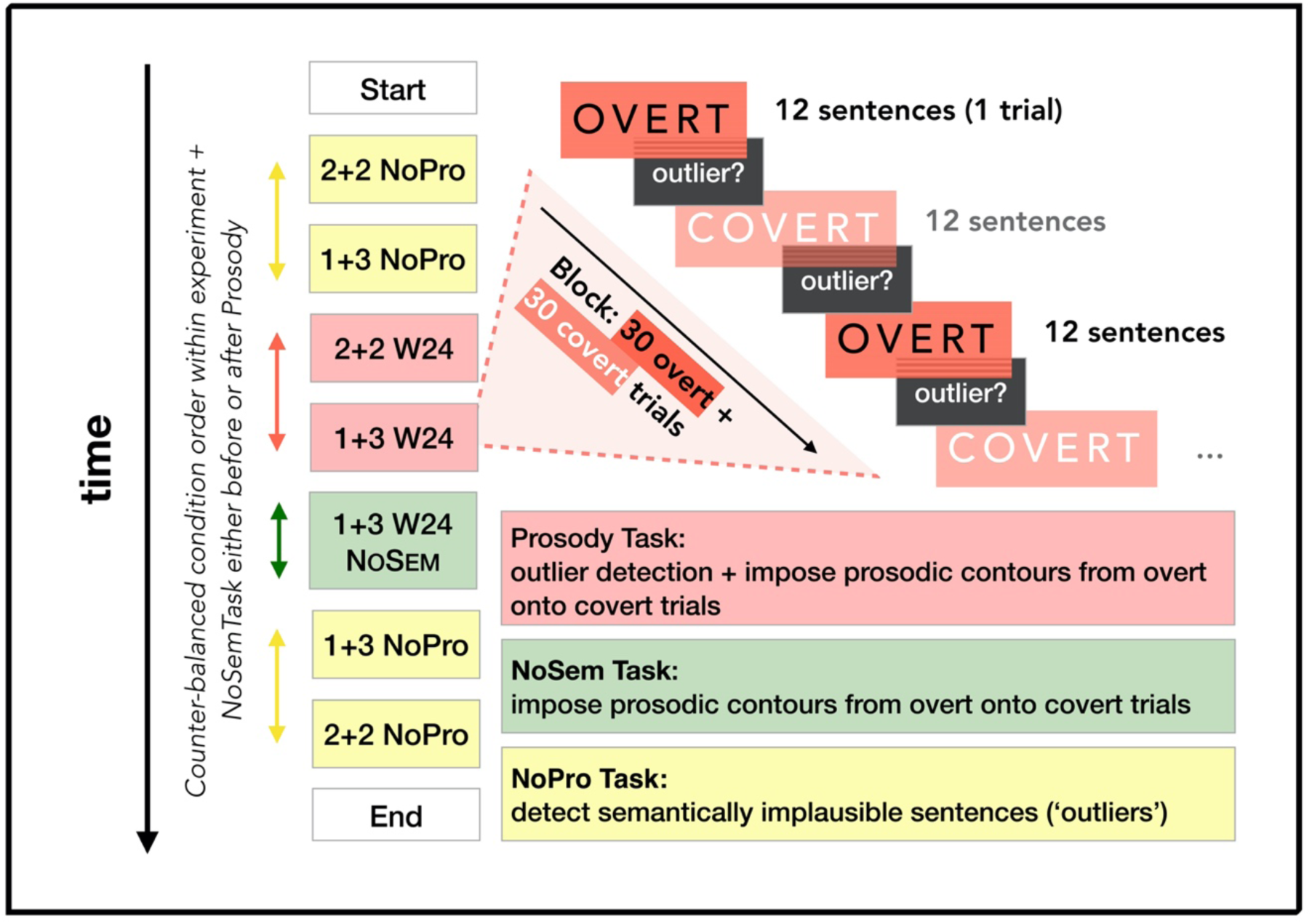
Experimental flow. Left: order of the three experiments and respective experimental blocks. In blue – the No Prosody experiments with the corresponding 1+3 and 2+2 Syntax blocks. In brown – the Prosody experiment. In green – the No Semantic Task, always represented by one block comprised of sentences with one of the syntactic structures. Right (top): scheme of experimental flow within one sample block of the Prosody experiment (2+2 W24 = 2+2 Syntax sentences with W24 prosodic contour). Starting with an overt trial (12 sentences), it continues with an outlier detection prompt, and then with the covert trial, during which participants listen to sentences with neutralized prosody while imposing onto them the intonational contour they attended to in the preceding overt trial. In the bottom right corner is the list of tasks used in the experiments.

In the **No Prosody experiment**, participants listened to 30 (22 without and 8 with semantic outliers) trials containing the sentences with neutralized prosody. At the end of each trial (i.e., after listening to 12 consecutive sentences), they had to indicate via button press if that trial contained a sentence that did not make sense (an ‘outlier’) or if all sentences were plausible. Trials consisting of the sentences with the same syntactic structure (1+3 or 2+2 Syntax) formed a block, with the order of blocks being counter-balanced across participants.

In the **Prosody experiment**, trials were presented in pairs. Participants first listened to a trial of 12 sentences, all of which had identical syntactic structures and the same overt prosodic contour (e.g., 1+3 Syntax with W24 contour). This *Overt Prosody* trial was immediately followed by a second *Covert Prosody* trial of 12 sentences, which still had the same syntactic structure as before (here: 1+3 Syntax) but lacked any prosodic contour (similar to the No Prosody experiment). During this second trial, participants were asked to silently ‘imagine’ the same prosodic pattern they had just heard during the overt prosody trial (here: W24). In other words, participants were instructed to process the sentences while imposing a *covert* prosodic contour (see Figure 6b). This *Covert Prosody* trial inherited its prosodic characterization from the preceding overt prosody trial (i.e., 2+2 syntax with *covert* W24 contour). Comparing the EEG signals of these trials to the No Prosody conditions should reveal the contribution of both overt and covert prosody to the elicitation of power peaks. After each trial (with overt or covert prosody) participants had to indicate by button press if that trial contained a semantic outlier sentence or not.

The structure of the **No Semantic Task experiment** was similar to the Prosody experiment. However, in this experiment, participants were only presented with trials that did not include semantic outliers, and they only went through one of the experimental conditions (pseudo-randomly selected for each participant: for example, Syntax 1+3, Prosodic Contour W24; see Figure 3). After the participant listened to the first trial, they would click ‘Next’ to start the second trial. This condition was introduced to investigate the effect of the semantic outlier detection on the EEG data in the Prosody experiment. We predicted that the data will be characterized less by the effects of syntax-prosody alignment and more by the independent processing of prosodic changes. Results and detailed discussion of this experiment are presented in Supplementary Materials *C*.

### EEG recording and processing

EEG data were recorded at a 500 Hz sampling rate using 64 cap-mounted electrodes (extended International 10-20 electrode organization System, Jasper, 1958; Waveguard^™^ original ANT Neuro EEG system), referenced online to the right mastoid. Matlab (Mathworks, 2011) and EEGLAB (version 14_1_0b; Delorme & Makeig, 2004) were used for EEG data preprocessing (the code is available upon request). Offline, we re-referenced the data to linked mastoids, removed bridged electrodes (the values were interpolated from the neighbouring electrodes after extracting epochs from the data), and performed resampling of the continuous datasets to 250 Hz. We filtered the data separately with a low- (20 Hz cut-off, filter order = 152) and a high-pass (0.2 Hz cut-off, filter-order = 2266) FIR filter using the Kaiser window (β = 5.65326). We removed eye movement artifacts using Independent Component Analysis (ICA; Lee, Girolami, & Sejnowski, 1999) run on the strongly high-pass filtered copies of the original datasets (1 Hz cut-off, filter order = 454). Note that we used these datasets for ICA decomposition only, for which we cut them into dummy epochs that underwent automatic artifact removal. Epochs were removed whenever the EEG at any time point (1) exceeded the threshold of |400| μV, (2) deviated at any of the electrodes from the mean amplitude at that electrode by 2 SDs, or (3) deviated at any of the electrodes from the mean of activity at all electrodes by 6 SDs. We copied the results of the ICA decomposition back onto the continuous data from which the activity of the components accounting for eye movements was then subtracted.

For frequency tagging analysis, the data were cut into 14.08-sec long epochs time-locked to the beginning of the second (rather than the first) sentence in each trial to avoid transient noise associated with the processing of the beginning of a given trial. Epochs containing signal crossing the |40| dB threshold in the 0-4 Hz frequency range were removed.

The mean of each epoch was subtracted from each data point in it, after which EEG was averaged across trials resulting in one average for each participant, electrode, and experimental condition. We calculated the evoked power assessed using the fast Fourier transform (FFT) of time-domain EEG responses averaged across trials (i.e., the FFT of the ERP, representative of the power of brain activity synchronized with the speech input) as well as inter-trial phase coherence (i.e., the coherence of phase angles across trials, which can change differently from the evoked power; ITPC). The resulting resolution of the frequency data was 0.071 Hz.

Due to the uneven distribution of noise across the frequency domain, the evoked power was further normalized by dividing the power at every frequency bin value in the spectrum by the mean of the response power at 14 neighbouring bins comprising 0.5 Hz prior as well as 0.5 Hz following that frequency bin (7 bins of 0.071 Hz on each side of the target one). The resulting data can be seen as the signal-to-noise (SNR) ratio of the EEG power across the frequency spectrum.

### Statistical analysis

The R code for the statistical analyses is available upon request. Behavioural binomial generalized mixed-effects models (lme4 package in R, Bates, Maechler, Bolker, & Walker, 2015) were built following forward-directed model comparison based on the Akaike Information Criterion (AIC). Relevant details of the final models are presented in the Results section. For the analysis of behavioural responses in the No Prosody and Prosody experiments, we fitted two generalized binomial models. The first one tackled the effect of Prosody (Overt, Covert, or No Prosody). The fixed effects tested for inclusion into this model were Prosody, Syntax (1+3 vs. 2+2), and Item Type (Correct vs. Outlier). Random intercepts for each participant and item were included by default, prior to model comparison. All fixed effects included in the best model were then tried as random slopes when appropriate while the model was converging successfully and was not reaching singularity. The build-up procedure for the second model testing the potential effect of alertness and familiarity on response accuracy was similar, but the only fixed effects tested for inclusion were Experiment Part (Beginning vs. End), Syntax, and Item Type. Additionally, d-prime values were calculated to form one of the predictors of the EEG data (as response accuracy has been shown to correlate with sentence-level EEG effects by Ding and coauthors, 2017).

In the analysis of EEG data, first, normalized EEG power and ITPC at the sentence (0.78 Hz) and the ½ sentence (1.56 Hz) frequencies was tested using bias-corrected and accelerated bootstrap tests (as implemented in the wBoot R package; Weiss, 2016) against the normalized power at the neighbouring frequencies (7 frequency bins on each side from the target frequency bin). This was done separately for each experimental block (see Figure 3). All p-values were Bonferroni-corrected for multiple comparisons. We extended this analysis by directly and systematically study the normalized EEG power and ITPC at the target frequencies, minus the noise at the 14 surrounding frequency bins, across different experimental conditions in the Prosody and the No Prosody experiments using two generalized linear models (one for EEG power, another one for ITPC). The models were fitted with inverse Gaussian distribution and an identity link function as their error terms were right-skewed. To allow for the use of the inverse Gaussian distribution, we added a minimal constant to both dependent variables shifting the values into the positive range. The independent variables included into the model were hypothesis-driven and comprised the following highest-level interactions and the embedded lower-level effects: Prosody × Syntax × Frequency (Sentence vs. ½ of a sentence), Anteriority (Frontal vs. Central vs. Posterior channels); Laterality (Left vs. Medial vs. Right channels); and Frequency × d-prime values. Potential side effects of familiarity and alertness of the participants were investigated by building additional models for normalized EEG power (hereafter, EEG power) and ITPC analysis. In these, only the data from the No Prosody experiment were used, and we included the following highest-level interactions as well as the embedded effects: Experiment Part (Beginning vs. End) × Syntax × Frequency × Anteriority; Laterality; and Frequency × d-prime values.

All linear models were visually checked for normality and homoscedasticity of the residuals. Residuals crossing the mean+|2.5*SD| threshold were removed when their distribution was not normal (resulting in the loss of 1% of observations). The absence of multicollinearity was ensured based on the condition number (Belsley et al., 1980, cited in Baayen, 2008) and the variance inflation factor (Craney & Surles, 2002). The post-hoc analysis of interactions was performed by comparing the least-squares means and their standard errors (as implemented in the lsmeans package; Lenth, 2016).

We additionally studied the relationships between our EEG effects (i.e., the ½ sentence and the sentence rate EEG responses within and between different conditions) using Pearson’s correlation to investigate our interpretations regarding the (dis)similarity of some findings (see Results).

## Results

### Performance on the behavioural task

On average, participants were 72.2% accurate in assessing the semantic acceptability of the sentences (comparable to results of Ding et al., 2017). Performance was higher on trials without outliers than on those with semantically implausible sentences (β = 1.498, SE = 0.335, p < .001). Participants performed slightly better at assessing acceptability of trials with covert prosody compared to trials in the No Prosody conditions, with no difference found between Overt prosody and the other two conditions (CovP vs. NoP: β = 0.194, SE = 0.079, p = .04). In addition, we found that acceptability of trials without outliers within the No Prosody experiment improved towards the end of the study (β = 0.515, SE = 0.153, p = .004). Syntactic structure of sentences did not significantly improve the models and was not included as a fixed effect.

### EEG results

#### No Prosody experiment

EEG power and ITPC values at sentence and ½ sentence frequencies in 1+3 and 2+2 Syntax were significantly larger than noise (all p < .001; Figure 4c-f). Based on previous research (Ding et al., 2016) and our own data (Supplementary Materials *B*), this aligns with the role of prosody in the elicitation of these EEG effects (see Figure 1). The sentence-level effects were in line with our predictions for both types of sentences. At the same time, according to the syntactic account, the peak in the EEG spectrum at the ½ sentence frequency would be expected only in the case of the 2+2 Syntax (corresponding to the phrase frequency). We therefore conjecture that at least the ½ sentence EEG peak in the 1+3 Syntax condition^1^ was likely induced by mechanisms other than syntactic processing, and specifically, in our view, by participants having placed a covert prosodic boundary in the middle of the 1+3 Syntax sentences. We further tested this hypothesis by investigating the correlations between the different EEG peaks in the No Prosody experiment and by comparing their scalp distributions. We hypothesized that if the ½ sentence EEG peak in the 1+3 Syntax condition is driven exclusively by prosody, (1) its size would vary differently across participants than the other EEG peaks that are at least partly influenced by syntactic processing, and (2) this modulation might have a distinct scalp distribution.

**Figure 4.**
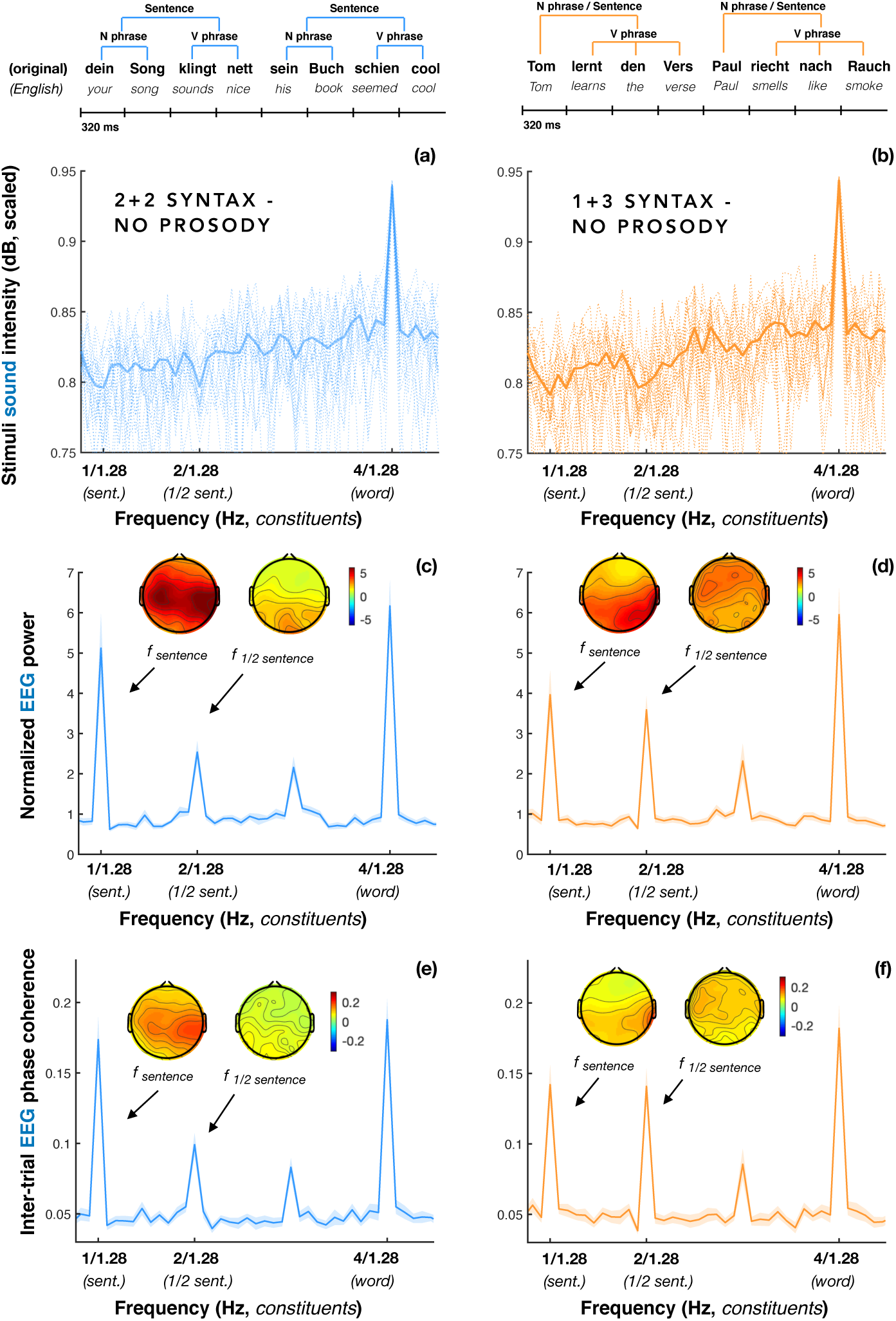
Spectral results: No Prosody experiment. Left panel (from top to bottom): sample sentences with the 2+2 Syntax structure, spectrum of sound intensity envelopes of the stimuli (panel *a*; thin dotted lines represent single trials, bold line depicts the average across all trials), the EEG power spectrum recorded while participants were listening to the 2+2 Syntax sentences (*c*), and the corresponding ITPC spectrum (*e*). Right panel: same for 1+3 Syntax. Note that the main syntactic boundary in the 1+3 Syntax condition (i.e., the one between first and second words) is not reflected in the spectrum, because the phrases forming the 1+3 Syntax condition are non-isochronous. The lines in the spectrum plots reflect group averages, with the shaded area depicting standard errors of the mean. Scalp maps depict scalp distribution of the EEG signal at the sentence and the ½ sentence frequencies (quantified as distance to the signal at surrounding frequencies). Note that the peak at 3/1.28 Hz (the rate at no syntactic or prosodic cues are modulated), represents a harmonic of the sentence frequency, similar to the 3 Hz response in the 1+3 Syntax condition in Ding and colleagues’ original study (2016).

Indeed, we found no correlation between the ½ sentence peaks in the two types of syntactic constructions (EEG magnitude: r^2^ = 0.033, p = .872; ITPC: r^2^ = -0.095, p = .645), while the sentence peaks were positively correlated (EEG magnitude: r^2^ = 0.479, p = .013; ITPC: r^2^ = 0.444, p = .023). The two ½ sentence peaks also had different scalp distributions: the effect in the 2+2 Syntax was centro-posterior (EEG power: frontal - posterior: β = – 1.544, SE = 0.271, p < .001; central - frontal: β = 0.768, SE = 0.214, p = .017), while in the 1+3 Syntax it was broadly distributed. Hence, it is plausible that the ½ sentence peaks in the two syntactic conditions were of different nature.

As the No Prosody experiment was run twice (once at the beginning and once at the end of the experiment), we compared these measures to estimate the effects of familiarity and alertness on results. While the EEG responses at the sentence and the ½ sentence frequencies were reduced at the end compared to the beginning of the experiment (EEG power: β = -1.025, SE = 0.081, p < .001; ITPC: β = -0.024, SE = 0.002, p < .001), this effect was not specific to the type of syntactic structure.

### Prosody experiment

The EEG data from the Prosody experiment are depicted in Figures 4 and 5. As in the case of the No Prosody sentences, spectral amplitude at sentence and ½ sentence frequencies in every experimental condition in the Prosody experiment was significantly larger than noise (all *p-values* < .001). We next analyzed the results by comparing peaks across experimental conditions to test our hypothesis about (i) the effects of overt and covert prosody as well as (ii) the syntax-prosody alignment on the EEG responses at frequencies of syntactic constituents. We found a significant effect of Prosodic Contour × Syntax in both power (β = 0.456, SE = 0.072, p < .001) and ITPC models (β = 0.015, SE = 0.002, *p* < .001). With respect to power, we also found a significant three-way interaction between Prosodic Contour, Syntax, and Frequency (β = -0.167, SE = 0.072, *p* = .02). That is, prosody had different effects on EEG spectral peaks elicited by the 2+2 compared to the 1+3 Syntax sentences.

**Figure 5.**
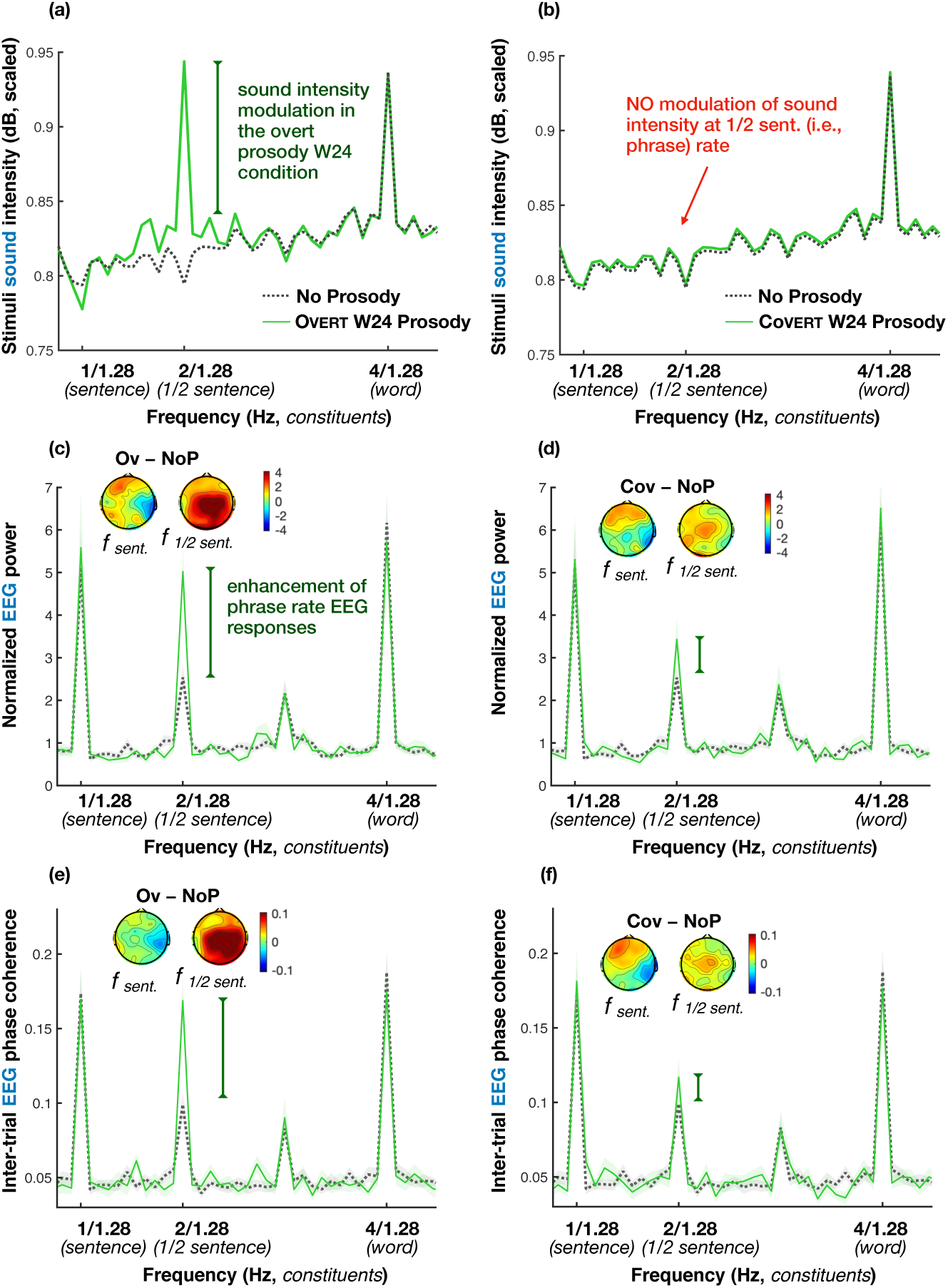
EEG results for the 2+2 Syntax sentences with the prosodic contour aligning with their syntactic structure plotted against the data from the same 2+2 Syntax sentences in the No Prosody experiment (dotted grey lines): overt prosody (left column) and covert prosody (right column) conditions. The top row (*a-b*) represents spectrum of sound intensity envelopes of the sentences, the middle and the bottom row depict EEG power (*c-d*) and ITPC spectra (*e-f*), respectively. The lines in the spectrum plots reflect group averages, with the shaded area depicting standard errors of the mean. Scalp maps show the scalp distribution of the difference between EEG peaks (calculated as distance from the peak value to surrounding noise) in the Prosody and the No Prosody experiments (separately for overt and covert prosody conditions and for sentence and ½ sentence frequencies). Key effects are marked with vertical lines (green for Prosody > No Prosody): when a prosodic contour is applied to the sentences in which it aligns with syntactic phrasing (whether overtly or covertly), the EEG responses were enhanced compared to the condition with no overt or instructed prosody (No Prosody).

**Figure 6.**
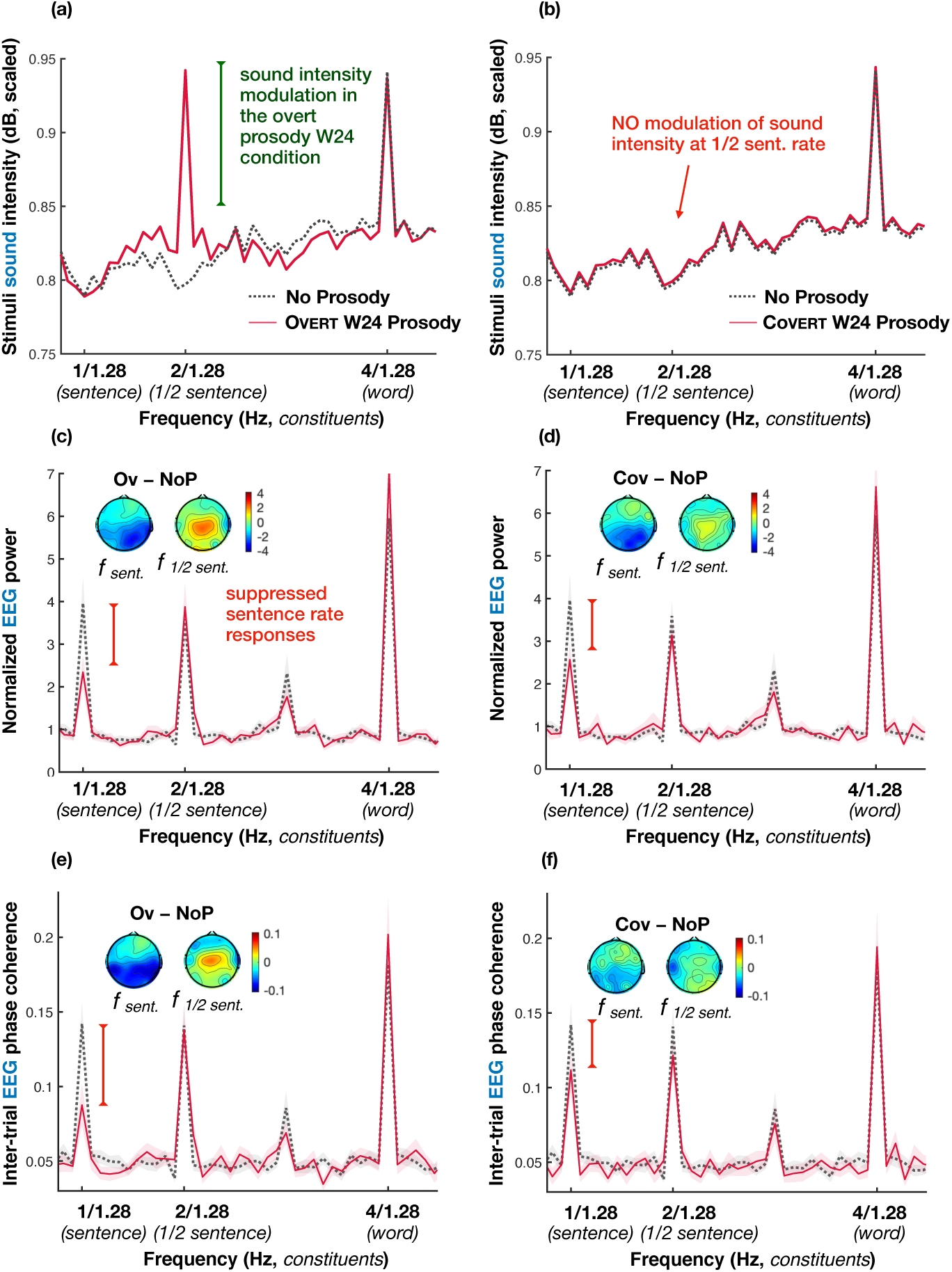
EEG results for the 1+3 Syntax sentences with the prosodic contour misaligned with their syntactic structure (thin red lines) plotted against the data from the same 1+3 Syntax sentences in the No Prosody experiment (dotted grey lines): both overt prosody (left column) and covert prosody (right column) conditions. The top row represents the spectrum of the sound intensity envelopes of the sentences (*a-b*), the middle and the bottom rows depict normalized EEG power (*c-d*) and ITPC spectra (*e-f*) respectively. The lines in the spectrum plots reflect group averages, with the shaded area depicting standard errors of the mean. Scalp maps show the scalp distribution of difference between EEG peaks (calculated as distance from the peak value to surrounding noise) in the Prosody and No Prosody experiments, separately for overt and covert prosody conditions and for the sentence and the ½ sentence frequencies. Prominent significant effects are marked with vertical lines (red for Prosody < No Prosody). When a prosodic contour not aligning with the syntactic structure was overtly or covertly applied to the sentences, EEG responses were diminished compared to the condition with no overt or instructed prosody.

Figure 5 illustrates the comparison between the No Prosody and the W24 prosodic contour in the case of syntax-prosody *alignment* – i.e., in 2+2 sentences, depicting first the sound intensity spectrum of the speech signals (panels a and b) and then the spectra for EEG signal (panels c-f). When participants listened to 2+2 Syntax sentences with an *overt* W24 prosodic contour (left panel), EEG power and ITPC were higher compared to the NoP but not compared to the CovP condition (EEG power: NoP-OvP: β = -1.126, SE = 0.223, p < .001; CovP-OvP: β = -0.678, SE = 0.259, p = .094; ITPC: NoP-OvP: β = -0.028, SE = 0.006, p < .001; CovP-OvP: β = -0.017, SE = 0.007, p = .188). EEG power enhancement was largely driven by the responses at the ‘supported’ by prosody ½ sentence frequency (NoP-OvP: β = - 2.156, SE = 0.29, p < .001).

Given the substantial impact the W24 pattern had on the ½ sentence peak in term of the acoustic spectrum (Figure 5a, first row), it could be argued that the corresponding EEG changes at this frequency (in rows 2 and 3) might, in principle, be driven by bottom-up acoustic changes. Crucially, however, the covert prosody condition resulted in similar EEG changes as the overt prosody condition. That is, when participants were presented with prosodically neutralized versions of the exact same 2+2 Syntax sentences - but were asked to simply imagine the aligned with the syntax W24 prosodic contour in absence of any prosodic cues in the speech signal (see Figure 5b, first row) –, we once again observed very similar EEG effects (Figure 5b, rows 2 and 3). Again, the ½ sentence EEG power was *larger* for the sentences with the *covert* W24 prosodic contour compared to the NoP condition (β = -0.82; SE = 0.211; p = .006).

In contrast to the 2+2 Syntax, when 1+3 Syntax sentences were presented with the overt W24 contour (i.e., the prosodic contour *misaligned* with the 1+3 Syntax structure), this combination elicited EEG power that was significantly *smaller* than in the NoP condition (EEG Power: NoP-OvP: β = 0.524, SE = 0.162, p = .016; ITPC: NoP-OvP: β = 0.032, SE = 0.005, p < .001). This effect of syntax-prosody misalignment was largely driven by the responses at the sentence rate (EEG power: β = 0.958, SE = 0.194, p < .001). Importantly, an analogous suppression of EEG responses was observed when the W24 prosodic contour was applied to the sentences covertly, or imagined by the participants (NoP-CovP: EEG power: β = 0.636, SE = 0.156, p < .001; ITPC: β = 0.021, SE = 0.005, p < .001; see Figure 5).

The effect of the W24 prosodic contour was as predicted. When prosody was aligned with syntactic structure, cortical responses at the syntactic constituent rates were enhanced compared to sentences with neutralized prosody. When, on the other hand, the prosodic contour did not align with syntactic phrasing, we saw reduced cortical responses compared to sentences with neutralized prosodic cues. These effects were specific to the frequency of syntactic constituents that were prosodically emphasized (in the case of the syntax-prosody alignment) or de-emphasized (in the case of a misaligned contour). Moreover, the effects of overt and covert prosody were found to be strikingly similar. This extends to the fact that participants with a larger enhancement of ½ sentence rate responses by the W24 contour overtly applied to 2+2 Syntax sentences were also the ones with the larger effect in the CovP condition. Similarly, larger suppression of the sentence-level responses by overt W24 prosody in the 1+3 Syntax sentences was associated with larger suppression in the CovP trials (see Figure 7).

**Figure 7.**
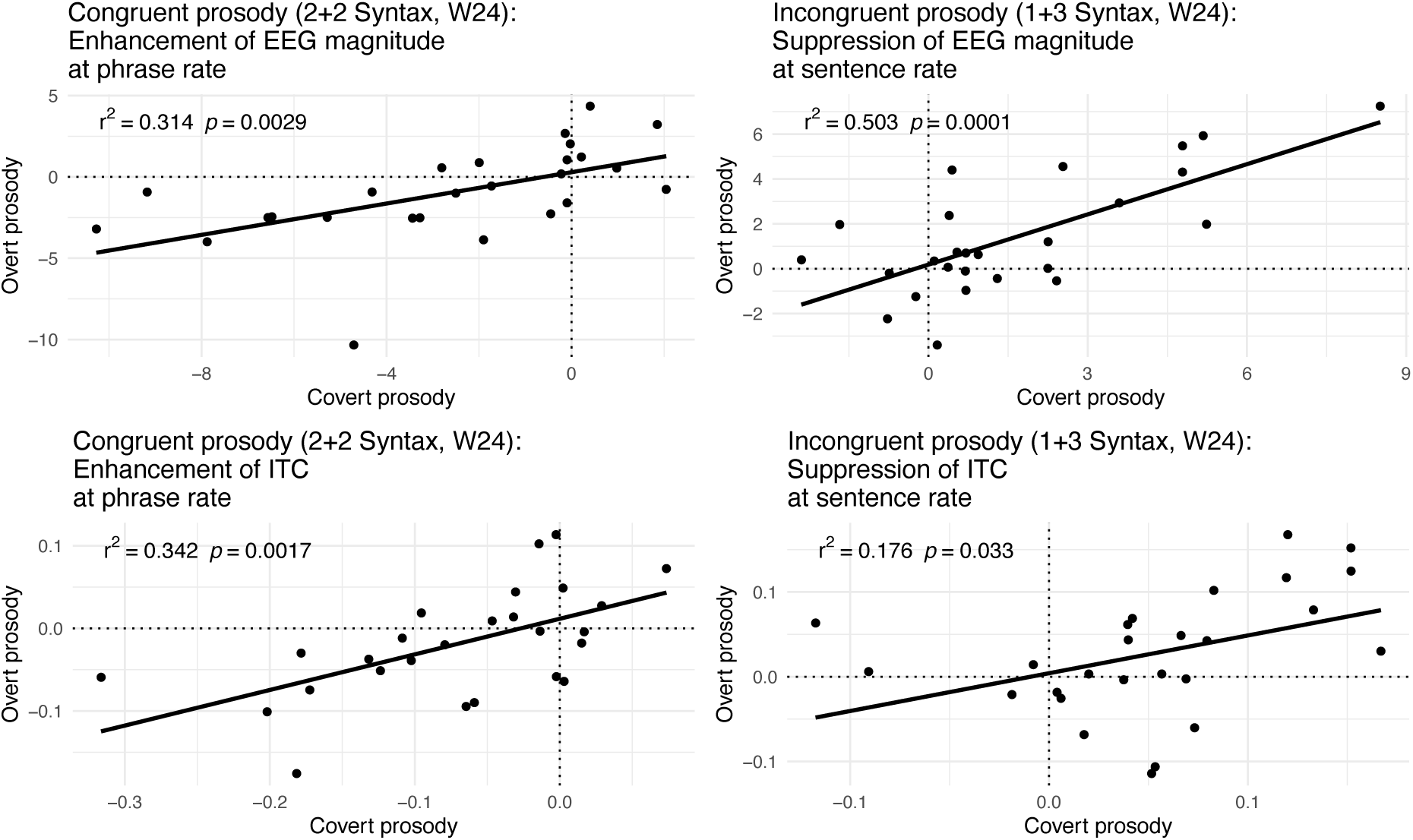
Correlations between effects of overt and covert W24 prosodic contour aligned with the 2+2 Syntax and misaligned with the 1+3 Syntax sentences. Left panel: the ½ (phrase) rate enhancement of EEG responses in the case of syntax-prosody alignment (2+2 Syntax W24 prosodic contour) contrasted with the No Prosody EEG spectrum peak. Right panel: the sentence rate suppression of EEG responses in the case of syntax-prosody misalignment (1+3 Syntax W24 prosodic contour). The values represent differences in normalized EEG power (in arbitrary units, scaled, top row) and IT(P)C (bottom row) between No Prosody and Prosody (NoP-OvP and NoP-CovP) experiments at either ½ sentence frequency (in the case of 2+2 Syntax sentences on the left) or sentence frequency (in the case of 1+3 Syntax sentences on the right). That is, at zero, there is no difference between NoP and the prosody condition; negative values reflect enhancement of the effect by the application of the W24 prosodic contour; positive values reflect reduction of the effect in the W24 compared to the NoP condition.

Finally, the relevance of the neurophysiological effects reported to the behavioural reality of sentence processing is emphasized by the fact that sentence-level EEG responses are positively associated with the performance on the behavioural task (d-prime: EEG power: β = 0.393, SE = 0.052, p < .001; ITPC: β = 0.012, SE = 0.002, p < .001; Frequency × d-prime: EEG power: β = 0.327, SE = 0.052, p < .001; ITPC: β = 0.007, SE = 0.002, p < .001), and similar to Ding and colleagues’ (2017) results, this association is much stronger than the one for the ½ sentence rate responses (for additional investigation of task effects, see results of our No Semantic Task experiment in Supplementary Materials *C*).

## Discussion

In running speech, we *hear* syntactic groupings typically only if they are expressed prosodically by boundary markers such as breaks, syllable lengthening, or pitch changes. However, prosodic groupings are not always driven by syntactic structure; prosodic boundaries can also be motivated by non-syntactic principles, such as the ‘same sister’ constraint that leads to prosodic phrases with equal numbers of syllables, independent of syntactic constituents (Fodor, 1998; Frazier, Carlson, & Clifton, 2006). Because syntactic and prosodic processing are distinct but closely associated, accurate understanding of sentence processing mechanisms requires careful consideration of both prosody and syntax and their interaction. Even when a speech signal is lacking overt prosodic cues (like in Ding et al., 2016), language users can and do imagine implicit, or covert, prosodic contours during processing (Fodor, 1998; 2002; Steinhauer, 2003). Over three experiments, we studied how neural responses to linguistic phrases and sentences are modulated by both overt and covert prosody. We used the frequency tagging approach, a method recently applied to the study of sentence processing. Although results from recent frequency tagging studies (Ding et al., 2016, 2017) have been interpreted as evidence for cortical tracking of hierarchical syntactic structure, our data show that other top-down mechanisms, and covert prosodic phrasing in particular, can account for most of these effects. We showed that low frequency cortical activity tracks both overt and covert prosodic changes, and this tracking interplays with syntactic processing.

We approached the investigation of prosodic processing in several steps. First, we conducted an EEG pilot study testing if *non-word groupings* based exclusively on pitch manipulations (2+2: low-low-high-high versus 1+3: low-high-high-high) would replicate Ding et al.’s MEG findings for 2+2 and 1+3 syntactic structures. As expected, we found a significant EEG power peak corresponding to a 2-word grouping (½ sequence peak in Supplementary Materials *B*) only for the 2+2 but not for 1+3 grouping condition.

Next, we studied real sentences with 2+2 and 1+3 phrasal groupings in which overt prosodic cues were neutralized (*No Prosody experiment*), much like in Ding and colleagues’ study (2016). However, unlike Ding and colleagues (2016), we unconfounded syntactic and possible covert prosodic structures such that 1+3 sentences were compatible with a prosodic boundary in mid-sentence position. Both EEG power and ITPC measures showed strong peaks at sentence and ½ sentence frequencies. A purely syntactic account can explain the large ½ sentence peaks for the 2+2, but not for the 1+3 structure – after all, Ding and colleagues (2016) used the absence of this peak in their 1+3 structures as evidence for a structural account reflecting syntactic phrase processing. Our tentative interpretation is that the ½ sentence peak in the 2+2 condition reflects top-down syntactic processing (in line with previous studies), whereas the corresponding peak in 1+3 sentences emerges from listeners’ spontaneous covert prosodic phrasing. This qualitative difference between sentence conditions was supported by both (a) a significantly more posterior scalp distribution of the ½ sentence peak in 2+2 than 1+3 sentences, and (b) the lack of a significant correlation between the peak amplitudes in the two conditions. The former finding suggests that cortical tracking of prosodic phrases (in 1+3 sentences) is associated with more frontal activity at the scalp than tracking of syntactic phrases (this interpretation is also in line with the task effects reported in in the No Semantic Task experiment; see Supplementary Materials *C*). These differences would not be expected if the peaks reflected the same cognitive processes in both conditions. Keeping in mind the absence of the ½ sequence peak in our pilot nonword experiment, it is implausible that a harmonic account could explain the ½ sentence peak. Covert prosody can, moreover, arguably account for some aspects of the recent frequency tagging results investigating the relationship between harmonic structure and sentence grammaticality (Tavano et al., 2020).

In contrast to the ½ sentence peaks, the power peaks at full sentence frequency were significantly correlated between syntactic structures in terms of amplitude, and their scalp distributions were found to be indistinguishable, thus suggesting similar underlying cognitive processes across 2+2 and 1+3 sentence structures. In absence of any acoustic markers for sentence boundaries in the speech signal, the power peaks at sentence frequency must reflect the top-down integration of four words into a coherent sentence representation. Whether this cognitive process is primarily syntactic in nature (as previously suggested by Ding and colleagues, 2016) or stems from semantic processing (Frank & Yang, 2018) or combinatorics (Martin and Doumas 2017) is not immediately evident. Importantly, covert prosodic processing can be driven by both semantic and syntactic cues (Itzhak et al, 2010; Fodor, Nickels, & Schott, 2018), meaning the sentence-level peak can be mediated by prosody as well.

Several lines of previous research support the idea that sentence prosody is tracked by slow neural activity. Studies of the Closure Positive Shift (CPS; Steinhauer, 2003) showed that both overt and covert prosodic boundary processing is reflected in a slow event-related potential (ERP) component lasting for up to 400-500 ms. At the same time, we know that the time window corresponding to the rate of cortical delta oscillations (1-4 Hz) encompasses the length of acoustic chunks that can be efficiently processed in behavioural paradigms (Ghitza, 2017; Rimmele et al. 2020). The link between the delta oscillatory range and behavioural data on the delta oscillatory range is supported by Meyer and colleagues (2017) reporting that delta oscillation phase changed at phrase boundary positions, whether phrase boundaries were driven by prosodic change or by parsing choices (in the absence of overt prosodic cues). While no frequency tagging studies to date have studied prosodic processing per se, it has been shown that imagined meter at 0.8 and 1.2 Hz is capable of eliciting significant EEG activity peaks in the absence of overt acoustic changes as well (Nozaradan, Peretz, Missal, & Mouraux, 2011). It is, therefore, highly plausible that a ½ sentence rate EEG peak in the 1+3 Syntax condition was driven by covert prosodic grouping.

Our *Prosody experiment* provided additional support for the role of prosody in the elicitation of low-frequency neurophysiological activity peaks. The overt prosody condition again elicited significant peaks at ½ sentence and sentence frequencies for both syntactic structures. While this result confirms that a prosodic contour is indeed sufficient to elicit the ½ sentence peak in absence of isochronous syntactic phrases (in 1+3 sentences), a concern is that this peak may be driven in a bottom-up fashion by acoustic cues already present in the speech signal (see Figures 4a and 5a). However, as expected, interactions between syntactic structure (2+2 vs 1+3) and prosodic contour (NoProsody vs W24) revealed that EEG activity peaks were differentially affected by the two prosodic contours. When the W24 contour aligned with syntax (in 2+2), the EEG was significantly enhanced (especially for the ½ sentence peak). When the contour was misaligned with syntactic phrasing (in 1+3), EEG peaks significantly decreased in amplitude (especially at the sentence frequency). This diverging pattern is clearly incompatible with a simple stimulus-driven account and points to higher cognitive processes that integrate both syntactic and prosodic information.

Data from our covert prosody conditions are further in line with this view. Recall that in these trials, participants listened to the same sentences with neutralized prosodic cues that were used in the No Prosody experiment. Similar to a covert prosody paradigm previously studied using event-related potentials (ERP) (Steinhauer & Friederici, 2001), participants had to silently imagine and superimpose the W24 prosodic contour they heard in the preceding overt prosody trial. While behavioural data demonstrated that participants still reliably identified semantic outliers, the EEG data clearly showed the effects of imagined, covert prosody. First of all, we again found the same elicited EEG peaks as the other experiments, including the ½ sentence peak for 1+3 structures. As no prosodic cues were present in the speech signal, this finding not only illustrates that participants were successful in silently imagining and imposing the W24 contour but also provides compelling evidence that the ½ sentence peak in 1+3 sentences can be elicited by a prosodic top-down mechanism, i.e., covert prosody. Intriguingly, the covert prosody conditions replicated the differential effects of the W24 contour that we previously observed for overt prosody. When aligned with the syntactic structure (in 2+2 sentences), the EEG activity peaks were enhanced, whereas they were decreased in the case of prosodic misalignment with syntactic phrasing in 1+3 structures. The effects of overt and covert prosody were tightly linked, as reflected in their significant correlation in both EEG power and ITPC. Similar parallels between overt and covert prosodic processing have been found in numerous psycho- and neurolinguistic studies (e.g., Hwang & Steinhauer, 2011; Itzhak et al., 2010; Kerkhofs, Vonk, Schriefers, & Schriefers, 2007; Drury, Baum, Valeriote, & Steinhauer, 2016; Steinhauer, 2003).

The experiments demonstrate the contribution of overt (old news) and covert (new news) prosody to low frequency cortical activity tracking sentence structure. While it is not out of the question that in addition to the prosodic effects, low frequency cortical activity tracks syntactic or semantic constituents (see results of our No Semantic Task experiment in Supplementary Materials *C*), the prosody-only account is the most parsimonious one. Whether syntactic structure is tracked by slow neural activity as well or if its processing is realized through other mechanisms (reflected, for example, in the high-gamma envelope changes; Nelson et al., 2017), is yet to be determined. Regardless of the frequencies of the neural activity tracking syntax and prosody in sentences, it is evident that the two mechanisms are interactive, and their integration is reflected in slow cortical responses.

## Data availability

The data that support the findings of this study as well as the R and Matlab scripts are available from the corresponding author upon request.

## Footnotes

1 The elicitation of this peak was also replicated in a larger group of participants (N=36) who only took part in the first run of the No Prosody experiment.

