## Supplementary Materials A, B, C for "Overt and covert prosody are reflected in neurophysiological responses previously attributed to grammatical processing"

MANUSCRIPT 1: SUPPLEMENTARY MATERIALS

SUPPLEMENTARY MATERIALS *A*

**Full list of 1+3 Syntax sentences (English translations in parenthesis)**

Max baut das Haus. (Max is building the house.)

Hein fährt nach Bonn. (Hein is going to Bonn.)

Karl fand den Grund. (Karl found the reason.)

Fritz fliegt nach Neuss. (Fritz is flying to Neuss.)

Franz fragt nach Tee. (Franz is asking for tea.)

Ben fuhr mit mir. (Ben went with me.)

Till geht nach Rom. (Till is going to Rome.)

Kurt geht zum Turm. (Kurt is walking to the Tower.)

Jan giert nach Lob. (Yann is craving for praise.)

Lex greift nach Karl. (Lex is reaching out for Karl.)

Chris kennt das Buch. (Chris knows the book.)

Dirk kommt nach Laos. (Dirk comes to Laos).

Bert läuft zum Tor. (Bert is running to the gate.)

Tom lernt den Vers. (Tom is learning the verse.)

Lars mag das Bild. (Lars likes the picture.)

Phil maß das Bad. (Phil was measuring the bathroom.)

Hans mied den Rum. (Hans was avoiding the rum.)

Tim muss nach Kiel. (Timm has to go to Kiel.)

Klaus plant das Fest. (Klaus is planning the party.)

Knut putzt das Klo. (Knut is cleaning the toilet.)

Nick reist nach Lund. (Nick is travelling to Lund.)

Alf rennt zum Fluss. (Alf is running to the river.)

Paul riecht nach Rauch. (Paul is smelling like smoke.)

Mark ringt nach Luft. (Mark is gasping for air.)

Gert roch den Gin. (Gert smelled the gin.)

Ron ruft nach Lars. (Ron is calling Lars.)

Rolf rührt den Teig. (Rolf is stirring the dough.)

Jörg ruht seit neun. (Jorg has been resting since nine.)

Joern sah zum Baum. (Jorn looked at the tree.)

Ken schleicht zum Sitz. (Ken is sneaking to the seat.)

Kai spart das Geld. (Kai is saving the money.)

Rick spricht zum Volk. (Rick is speaking to the people.)

Sven stinkt nach Schweiss. (Sven stinks of sweat.)

Horst strebt nach Glück. (Horst is striving for happiness.)

Jens sucht das Mehl. (Jens is looking for flour.)

Nils wäscht das Hemd. (Nils is washing the shirt.)

Ralf wies zum Rad. (Rail is pointing to the bike.)

Kim wischt den Flur. (Kim is wiping the hallway.)

Wim zahlt das Bier. (Wim is paying for the beer.)

Phil zieht nach Prag. (Phil is moving to Prague.).

**Full list of 2+2 Syntax sentences**

Der Bär trinkt erst. (The bear is drinking first.)

Mein Baum steht da. (My tree is [standing] there.)

Das Bett riecht frisch. (The bed smells fresh.)

Das Blatt fliegt her. (The leaf is flying here.)

Mein Boot kippt um. (My boat tips over.)

Mein Boss spricht klar. (My boss speaks clearly.)

Dein Brot schmeckt schlecht. (Your bread tastes bad.)

Sein Buch schien cool. (His book seemed cool.)

Der Busch wächst schnell. (The bush is growing quickly.)

Ihr Chor sang schlimm. (Her choir sang poorly.)

Der Clown lacht lahm. (The clown is laughing lamely.)

Der Dieb lügt krass. (The thief is lying blatantly.)

Sein Fall lief schief. (His case went wrong.)

Sein Feld blüht schön. (His field is blossoming beautifully.)

Der Fisch schwimmt flink. (The fish is swimming nimbly.)

Mein Freund reist gern. (My friend likes to travel.)

Dein Holz brennt schnell. (Your wood burns fast.)

Sein Hund hört nicht. (His dog does not listen.)

Ihr Kerl spielt falsch. (Her guy is playing incorrectly.)

Ihr Kind schläft dort. (Her child is sleeping there.)

Ihr Kleid ist naß. (Her dress is wet.)

Ihr Koch backt nie. (Her cook never bakes.)

Mein Kopf juckt rechts. (My head is itching on the right.)

Sein Licht scheint schwach. (His light seems weak.)

Ihr Mann sieht kaum. (Her husband barely sees.)

Das Paar wohnt links. (The couple lives on the left.)

Sein Pferd trabt schwer. (His horse is trotting heavily.)

Mein Pool roch fies. (My pool smelled nasty.)

Das Reh fällt hin. (The is falling.)

Das Schaf war naß. (The sheep was wet.)

Dein Schal weht rum. (Your scarf is blowing around.)

Der Schnee schmilzt früh. (The snow metls early.)

Dein Schrank wog viel. (Your dresser weighed a lot.)

Dein Sohn ritt fort. (Your son rode away.)

Dein Song klingt nett. (Your song sounds nice.)

Das Team malt selbst. (The team paints itself.)

Mein Vieh ging ein. (My kettle is dying off.)

Der Wolf biss oft. (The wolf bit often.)

Das Zelt blieb hier. (The tent stayed here.)

Sein Ziel hing hoch. (His goal was [hung] high.)

**Outlier sentences: 1+3 Syntax**

Max baut den Rauch. (Max is building the smoke.)

Rolf rührt das Haus. (Rolf is mixing the house.)

Jan giert nach Kiel. (Yann is craving Kiel.)

Lex greift nach Prag. (Lex is attacking Prague.)

Dirk kommt nach Tee. (Dirk is coming to tea.)

Kim wischt das Glück. (Kim is washing happiness.)

Nils wäscht das Mehl. (Nils is washing the flour.)

Hein fährt nach Grund. (Hein is going to the reason.)

Knut putzt den Vers. (Knut is cleaning the verse.)

Kurt geht nach Lob. (Kurt is going to the praise.)

Gert roch das Fest. (Gert smelled the celebration.)

Tom lernt das Hemd. (Tom is learning the shirt.)

**Outlier sentences: 2+2 Syntax**

Dein Brot sang schlimm. (Your bread sang poorly.)

Der Bär blüht schön. (The bear is blossoming beautifully.)

Mein Boot trinkt erst. (My boat is drinking first.)

Dein Schal trabt schwer. (Your scarf is trotting heavily.)

Das Zelt lacht lahm. (The tent is laughing lamely.)

Der Busch schläft dort. (The bush is sleeping there.)

Dein Holz lügt krass. (Your wood is lying blatantly.)

Sein Fall biss oft. (His fall bit often).

Der Wolf schmilzt früh. (The wolf melts early.)

Sein Feld wohnt links. (His field lives on the left.)

Sein Licht juckt rechts. (His light is itching on the right.)

Das Bett spielt falsch. (The bed is playing incorrectly.)

**Stimuli intelligibility across OvP conditions and for the NoP sentences**

Participants were presented with all 104 sentences one-by-one. They could listen to every sentence maximum twice and had to type in what they understood. Response was considered correct if the sentence typed in by the participant was identical to the one presented, or if there were minor orthographical error and/or if one proper name was mistaken by another one while not changing the meaning of the sentence. The EEG (and behavioural) data from the participant who scored 49 out of 104 on the NoP sentences were not analyzed. Note that the slight difference in ranking between NoP and OvP sentences could be caused by the fact that participants in the OvP intelligibility pilot study were recruited separately from the main study and did not get to read the full list of sentences prior to the sentence intelligibility test. Thirty-seven participants performed the NoP sentence intelligibility task, however, only 25 of them went through the full version of the subsequent EEG study.

(1) Sentence intelligibility: NoP

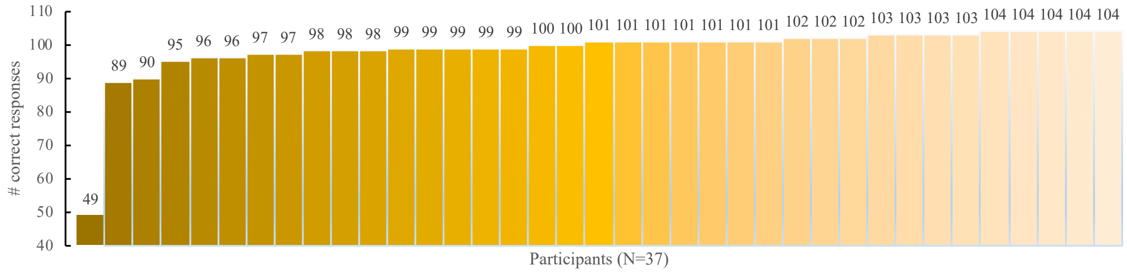


(2) Sentence intelligibility: W24

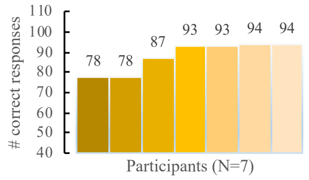

SUPPLEMENTARY MATERIALS *B*

**Control experiment: Non-syntactic grouping**

Ding and colleagues (2016) investigated the processing of 1+3 Syntax structures (for example, $fry_1$ | $to\text{-}ma\text{-}to_3$; see main text for details) using frequency tagging with MEG. To ensure that the results from this condition will be comparable to their data when EEG is used, we conducted a control experiment using non-syntactic rhythmic grouping. The reason for using non-syntactic grouping is that with natural language materials, many layers of information (e.g., semantics, syntax, prosody) can contribute to chunking of words into larger constituents. In our case, only one type of manipulation (rhythmic grouping differences) was ensured by using nonword materials. We compared processing of 2+2 and 1+3 grouping created by manipulating the average fundamental frequency at which the nonword was presented, resulting in artificially "sung" stimuli. Sequences made of four monosyllabic nonwords each contained two chunks: one presented with the average pitch of 123.47 Hz, corresponding to the B of the second octave, and the other one with the pitch of 164.81 Hz, corresponding to the E of the third octave (see below).

***Methods***

*Participants.* Eleven German native speakers participated in the experiment (age range: 20-35 years, mean age = 26; 6 women, 5 men). Participant inclusion criteria were identical to the ones in the main study.

*Materials.* We recorded a German native speaker pronouncing fifteen monosyllabic non-words that were phonotactically correct in German. The pitch in them was flattened, and the intensity was normalized in Praat (Boersma & Weenink, 2019) to 70 dB. The length of each nonword was manually adjusted to 250 ms. The nonwords were concatenated into sequences of four in a pseudorandom manner: within each sequence, no vowel or consonant was repeated, and no real words were created through concatenation. The 4-nonword sequences were concatenated into trials comprising 12 sequences each (48 nonwords). Each sequence lasted for one second, making trials 12 seconds long. No pauses were introduced at any time point within trials. Given the length of each nonword in this experiment was 250 ms, the nonword frequency corresponded to 4 Hz, the ½ sequence frequency to 2 Hz, and the sequence frequency to 1 Hz.

For each 4-nonword sequence, pitch was manipulated in a way specific to either the 1+3 or the 2+2 grouping. In the case of the 1+3 grouping, the first nonword was assigned a pitch of 123.47 Hz, and nonwords 2, 3, and 4 were assigned a pitch of 164.81 Hz. In the 2+2 grouping, the pitch of 123.47 Hz was imposed in Praat on the first two nonwords, and the pitch of 164.82 on nonwords 3 and 4. Note that sound intensity was controlled for, and sound intensity fluctuations were only observed at the nonword (i.e., the syllabic) rate (see left panel of Figure B1 in Supplementary Materials B).

To keep participants' attention, similar to the main study, we used an outlier detection task. Thirty trials per grouping were presented to each participant. In twenty two of these trials, all nonword sequences had the same (either 1+3 or 2+2) grouping. The 8 trials contained rhythmic outliers: either a 1+3 grouping sequence was inserted in a trial of 2+2 grouping sequences, and vice versa.

*Procedure.* This experiment was part of a series of pilot tests administered with the same group of participants. In total, participants came to the lab for approximately 4 hours. After filling out behavioural questionnaires (identical to the ones in the main study), participants were brought into the EEG booth, with the EEG session lasting for approximately 2 hours with several major breaks between tests. Participants were told that the sequences of syllables would form patterns. After each trial they were asked to respond if the main pattern within a trial has ever been broken or not. Trials were presented in blocks of 30, with blocks containing correct and outlier trials for one type of grouping (either 1+3 or 2+2).

*EEG recording and processing.* See main study for details. In contrast to the main study, single bin values in this experiment were calculated at the frequency of 0.0833 Hz (different from the main study due to the sequences of nonwords being shorter in this experiment).

*Statistical analysis*. We computed the average accuracy and the d-prime for the behavioural data in this pilot experiment in R (R Core Team, 2018). The EEG Magnitude was analyzed separately for the 1+3 and the 2+2 grouping using one-tailed paired t-tests contrasting the response at either the ½ sequence or the sequence frequency with the noise (average EEG magnitude at neighbouring bins comprising 0.5 Hz prior as well as 0.5 Hz following the target frequency). The p-values were corrected for multiple comparisons using the Bonferroni method.

***Results and discussion***

The accuracy on the outlier detection task was 66.1%, with the mean d-prime of 1.175 meaning participants did distinguish between the two types of rhythmic grouping. The EEG findings were in line with the behavioural data. The EEG responses at the sequence rate (1 Hz) were significantly different from noise in both 1+3 and 2+2 grouping types ($t = -6.169$, $p < .001$ and $t = -4.562$, $p = .001$, respectively. At the ½ sequence rate, as predicted based on the MEG results in Ding and colleagues' study (2016), the EEG magnitude was only significant from noise in the 2+2 grouping ($t = -3.106$, $p = .011$). These results suggest that 1+3 grouping can be used in EEG frequency tagging studies as a control condition for constructions with 2+2 grouping.

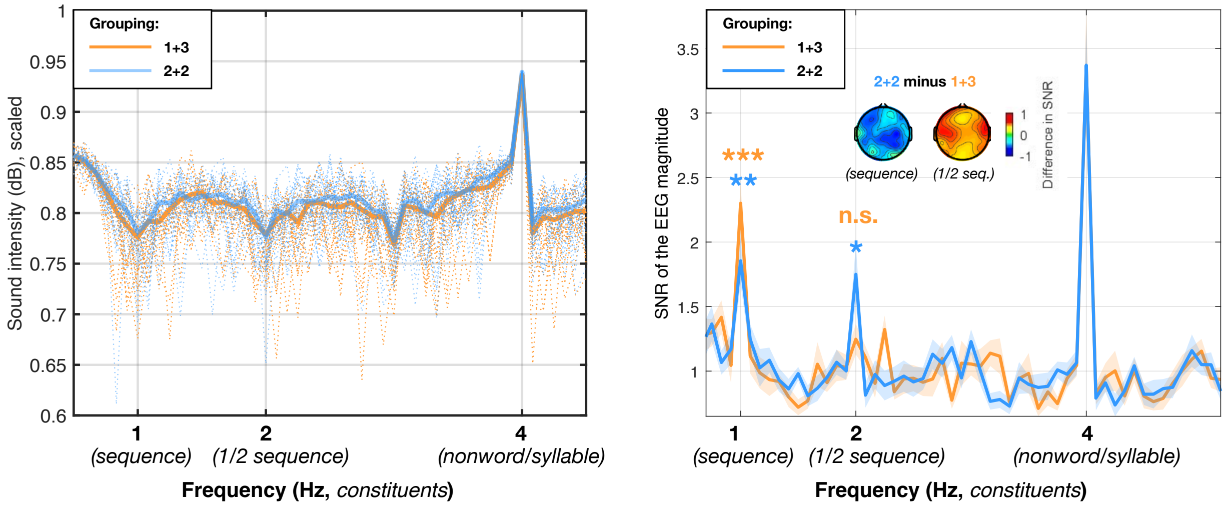


**Figure B1. Sound intensity envelope and the EEG magnitude spectra for 1+3 and 2+2 nonword grouping based on pitch changes ("sung" groupings).** On the left is the sound intensity envelope calculated per trial, converted to a logarithmic decibel scale, and scaled from 0 to 1. The envelope was computed for each trial separately (thin dotted lines) and then averaged across trials (thick solid lines). On the right is the EEG magnitude spectrum from 11 participants in this experiment. Note that while the EEG responses at the frequency of a full sequence are significantly greater than noise in both conditions, while at the ½ sequence frequency this is only the case for the 2+2 grouping. The scalp maps depict distribution of the differences between EEG magnitude SNR in the two conditions (2+2 minus 1+3 grouping) across electrodes, separately for the sequence and the ½ sequence frequency. $^{*}p<0.05$ $^{**}p<0.01$ $^{***}p<0.001$.

#### *References*

Ding, N., Melloni, L., Zhang, H., Tian, X., & Poeppel, D. (2016). Cortical tracking of hierarchical linguistic structures in connected speech. *Nature neuroscience*, *19*(1), 158.

R Core Team (2018). R: A language and environment for statistical computing. R Foundation for Statistical Computing, Vienna, Austria. URL https://www.R-project.org/.

### No Semantic Task experiment

Similarly to the Prosody experiment, participants listened to OvP and CovP trials, however, in the No Semantic Task experiment, outlier trials were excluded. Participants' task was to listen the OvP trial passively, and then listen to the CovP trial while imagining the intonation present in the preceding OvP sentences.

#### *Methods*

For the No Semantic Task experiments, we analyzed a subset of 10 participants who were presented with the prosodic contour investigated in the frame of the current study and had sufficient artifact-free EEG data (age range: 20-37 years, mean age = 26, age SD = 6; 5 females, 5 males). The data were analyzed for the 1+3 Syntax W24 (N=10) condition only (the one for which the data of at least 10 participants were available). The EEG effects in the No Semantic Task were contrasted with the corresponding conditions in the Prosody experiment. Two generalized linear models (inverse Gaussian family with identity link) models were fitted: one for normalized EEG power and another one for ITPC. In both of them we included the following interactions with respective lower-level effects: Task (Dual task vs. No Semantic Task) × Prosody (Overt vs. Covert) × Frequency; Laterality × Anteriority; and Task × Frequency × Anteriority. The rest of the methodological aspects of the experiment were identical to those in the Prosody experiment.

#### *Results and discussion*

We used the No Semantic Task experiment to investigate the effect of task-driven processing on the results of the Prosody experiment. We presumed that the EEG responses in the No Semantic Task experiment would reflect the shift of attention from semantic information to prosody, and this would be especially prominent in the CoP condition where participants had to actively perform a prosodic task. We analyzed the data from the 1+3 Syntax sentences and found that the signal at ½ sentence frequency was significantly larger than noise in the overt and covert W24 condition (EEG power and ITPC: all *p-values* $< .001$). This was also the case for the sentence frequency (EEG power: OvP: $p = .001$; CovP: $p = .003$; ITPC: OvP: $p =.028$; CovP: $p = .031$). Note, however, that in this sample of participants, the ITPC in the OvP condition of the Prosody experiment was not significantly larger than noise ($p = .257$).

In the 1+3 Syntax sentences where prosodic structure did not support syntactic phrasing, the absence of any semantic task enhanced EEG responses across ½ sentence and frequencies were enhanced (main effect of Task on EEG power: $\beta = 0.288$, SE = 0.102, $p = .005$). This effect was more prominent at sentence frequency in OvP ($\beta = 0.798$, SE = 0.262, $p = .047$) and ½ sentence frequency in CovP ($\beta = 1.408$, SE = 0.459, $p = .045$). As seen from Figure C1, at the sentence rate, in the CovP condition, we found that the EEG power at the sentence frequency was, in contrast, smaller in No Semantic Task compared to the Prosody experiment ($\beta = -0.809$, SE = 0.257, $p = .038$).

The scalp distribution differences between the No Semantic Task experiment and the Prosody experiment were in line with our findings in the No Prosody experiment. Task × Anteriority interaction (EEG power: $\beta = -0.315$, SE = 0.144, $p = .029$; ITPC: $\beta = 0.\ 11$, SE = 0.004, $p = .008$) reflected a that while No Semantic Task effects were broadly distributed (similar to the presumably ‘prosodic’ ½ sentence peak in 1+3 Syntax NoP sentences), the effects in the Prosody experiment were less prominent at frontal electrodes (similar to the more ‘syntactic’ ½ sentence peak in 2+2 Syntax NoP sentences).

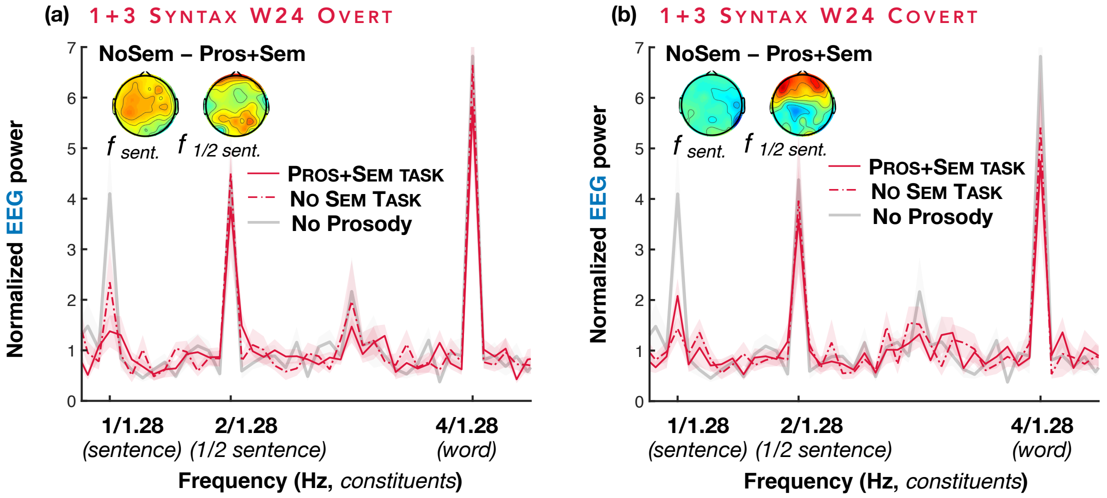


**Figure C1. EEG power in the No Semantic Task experiment: 1+3 Syntax sentences with overt (a) and covert (b) W24 prosodic contour.** Data from 10 participants per condition are plotted against the data from the same participants in the No Semantic Task (dashed-dotted red lines), the Prosody (solid red lines), and the No Prosody experiments (bold grey lines). The No Prosody condition is added to the graph to demonstrate the absence of the qualitative differences between the data of the subset of participants and the data in the main No Prosody experiment (however, the No Prosody condition was not included in the statistical analysis, for which only the comparison between the Prosody and the No Semantic Task experiment was quantified). The lines in the spectrum plots reflect group averages of the EEG magnitude SNR in the frequency domain, with the shaded area depicting standard errors of the mean. Scalp maps represent the scalp distribution of difference between the effects in the No Semantic Task and Prosody experiments.

We predicted that, without any explicit need to attend to sentence meaning, participants would shift their focus to the prosodic pattern, especially in the covert prosody trials where prosodic patterns must be actively generated via top-down mechanisms. We expected that effects related to prosody would be enhanced, and those related to semantic sentence integration reduced. What we found was an increased ½ sentence peak amplitude in both overt and covert prosody conditions. In line with our previous finding that prosody-driven ½ sentence peaks (in 1+3 sentences) tend to have a more frontal distribution than syntax-driven ½ sentence peaks (in 2+2 sentences), the increase of this peak had a similar scalp distribution extending to frontal channels, pointing to a stronger focus on prosodic processing when no semantic task was required. The sentence-rate EEG peak, in contrast, was affected by the removal of the semantic task differently between overt and covert prosody conditions, decreasing in covert prosody trials but slightly increasing in overt prosody trials. Since the positive correlation between sentence peak amplitude and semantic task performance directly pointed to its involvement in semantic processing, the decrease of this peak in absence of the semantic task was expected, especially in covert prosody trials (i.e., when subjects actively engaged in a prosodic mapping task). Sentence-level EEG activity enhancement in overt prosody trials, on the other hand, is somewhat difficult to interpret, as this is the only condition of our entire study where subjects did not perform in any task.
